# Proteomic profiling of IDH-wildtype Glioblastoma Tissue and Serum uncovers prognostic Subtypes and Marker Candidates

**DOI:** 10.1101/2024.02.29.582688

**Authors:** Tilman Werner, Agnes Schäfer, Michael Hennes, Miguel Cosenza Contreras, Guadalupe Espadas, Eduard Sabido, Lena Cook, Axel Pagenstecher, Niko Pinter, Tobias Feilen, Alexander Grote, Christopher Nimsky, Jörg Walter Bartsch, Oliver Schilling

**Affiliations:** Institute for Surgical Pathology, Faculty of Medicine, University Medical Center Freiburg; Faculty of Biology, University of Freiburg; Spemann Graduate School of Biology and Medicine (SGBM), University of Freiburg; MeInBio Graduate School, University of Freiburg, Freiburg, Germany; Department of Neurosurgery, Philipps University Marburg, Marburg, Germany; Proteomics Unit, Centre for Genomic Regulation (CRG), The Barcelona Institute for Science and Technology, Universitat Pompeu Fabra, Barcelona, Spain; Department of Neuropathology, Philipps University Marburg, Marburg, Germany; German Cancer Consortium (DKTK) and German Cancer Research Center (DKFZ), Heidelberg, Germany

## Abstract

**Background:** IDH-wildtype glioblastoma (GBM) is the most prevalent primary brain cancer with a 5-year survival rate below 10%. Despite combined treatment through extensive resection and radiochemotherapy, nine out of ten patients develop recurrences. The lack of targeted treatment options and reliable diagnostic markers for recurrent tumors remain major challenges.

**Methods & Aims:** In this study, we present the proteomic characterization of tissue and serum from 55 initial GBM tumors and five matching recurrences, which we investigated for proteomic tumor subtypes and proteomic signatures associated with recurrence.

**Results:** Primary tumors revealed four distinct subgroups through hierarchical clustering: a neuronal cluster with elevated mature neuron markers, an innate immunity cluster with increased protease expression, a mixed cluster, and a stem-cell cluster. Neurodevelopmental and inflammatory processes were identified as key factors influencing clustering, with proteolytic activity increasing relative to the degree of inflammation. An analysis comprising proteins with lower coverage confirmed and expanded this pattern. Patients in the neuronal cluster exhibited significantly longer survival compared to those in the stem-cell cluster. In a patient-matched differential expression analysis, five recurrent tumors displayed significantly altered protein expression compared to their primary counterparts, emphasizing the proteomic plasticity of recurrent tumors. Investigation of serum proteomes before and after surgery, using a depletion-based protocol, revealed highly patient-specific and stable proteome compositions, despite a notable increase in inflammation markers post-surgery. However, the levels of circulating proteolytic products matched to the proteolytic activity within the tissue and one fragment of proteolysis activated receptor 2 (PAR2) consistently dropped in abundance after removal of inflamed tumors.

**Conclusion:** Overall, we describe a large proteomic GBM cohort. We identified distinct tumor subgroups, molecular patterns of recurrence, and matching proteomic patterns in the bloodstream, which may improve risk prediction for recurrent GBM.

## Introduction

Isocitrate Dehydrogenase (IDH)-wildtype Glioblastoma (GBM) – formerly known as Glioblastoma Multiforme – is the most prevalent primary malignancy of the brain ^1,2^. It is also one of the deadliest cancers, with an average survival of 15-18 months and 5-year survival rates below 10% ^3,4^.

Although GBMs grow fast and are highly infiltrative, symptoms are variable and largely dependent on the affected brain area. Hence, they are usually detected rather late through Magnetic Resonance Imaging (MRI) or Computer Tomography (CT) ^2,5–7^. According to the World Health Organization (WHO) classification of brain tumors, GBM diagnosis is established, if an intracranial neoplasia presents an IDH- and Histone H3 wildtype in combination with at least one other qualifier like microvascular proliferation, necrosis, *TERT* promoter mutation, *EGFR* gene amplification or +7/-10 chromosome copy number changes. Due to their aggressiveness, all GBM are considered to be grade 4 tumors by the WHO, regardless of individual molecular or histological characteristics ^1^. So far, the etiology of GBM has remained unclear and only few risk factors are known ^1,8^. After diagnosis, patients usually undergo immediate surgical tumor resection in combination with chemoradiotherapy with Temozolomide (TMZ) ^2,9^. Nonetheless, the tumors return in virtually all cases, since infiltrative tumor cells can spread far beyond the surgical and radiotherapeutic margin into areas where a still intact blood-brain barrier shields them from chemotherapy ^1,10^. In addition, GBM is known for a remarkable cellular heterogeneity and genomic plasticity leading to drug resistances and even aggressive therapies can so far only be considered as life prolonging ^4,11,12^. Precise and effective diagnostic tools for recurrent GBM remain a major challenge. MRI cannot confidently detect recurrences at initial stages and is performed only infrequently (3-month intervals at best) ^9^. Therefore, more reliable and cost-effective diagnostic methods are crucial to complement the current diagnostic workup ^2^.

Current technologies such as explorative, unbiased, omics-type characterizations are ideally suited for identifying marker signatures and potential therapeutic targets ^13^. Different molecular subtypes of GBM have already been described in genomic and transcriptomic screens ^14,15^. Two proteogenomic studies by Wang et al and Yanovich-Arad et al adopted and refined these classifications into different subtypes with high neuronal activity and synaptic transmission, immune infiltration and extracellular matrix reorganization, or increased mRNA translation ^16,17^. Likewise, on a single-cell level, Neftel et al found similar patterns in a transcriptomic dataset and defined four cellular states of GBM development. Interestingly, although one cell state was usually dominant, cells in various states were present in all samples ^11^. The importance of neurodevelopmental, hypoxic, and inflammatory processes for the initiation of different transcriptional programs was also emphasized by Ravi et al in a spatial transcriptomic study ^18^. Across most publications, mutations and/or expression differences in *EGFR*, *PDGFRA*, and *NF1*, were highlighted as subtype-defining ^11,14–16^. Nevertheless, larger-scale investigations of GBM are needed - especially on the proteomic level, since mRNA expression is only loosely correlated with protein abundance ^19^.

Here, we present the proteomic characterization of 55 primary GBM tumors, five matching recurrences, and matching serum samples collected before and after each surgery.

## Materials and Methods

### Ethics and Sample Collection

All patients included in this study were informed and provided written consent. The study was approved by the local ethics committee (Marburg University, Medical Faculty) under file 185/12. All tissue samples were fresh-frozen in liquid nitrogen directly within the operation room and then stored at -80°C. Blood samples were collected one day before and three days after surgery in serum collection tubes. After centrifugation at 2000G for 10 minutes, serum samples were stored at -80°C. All included patients were diagnosed with IDH wildtype and WHO grade 4 GBM.

### Chemicals list

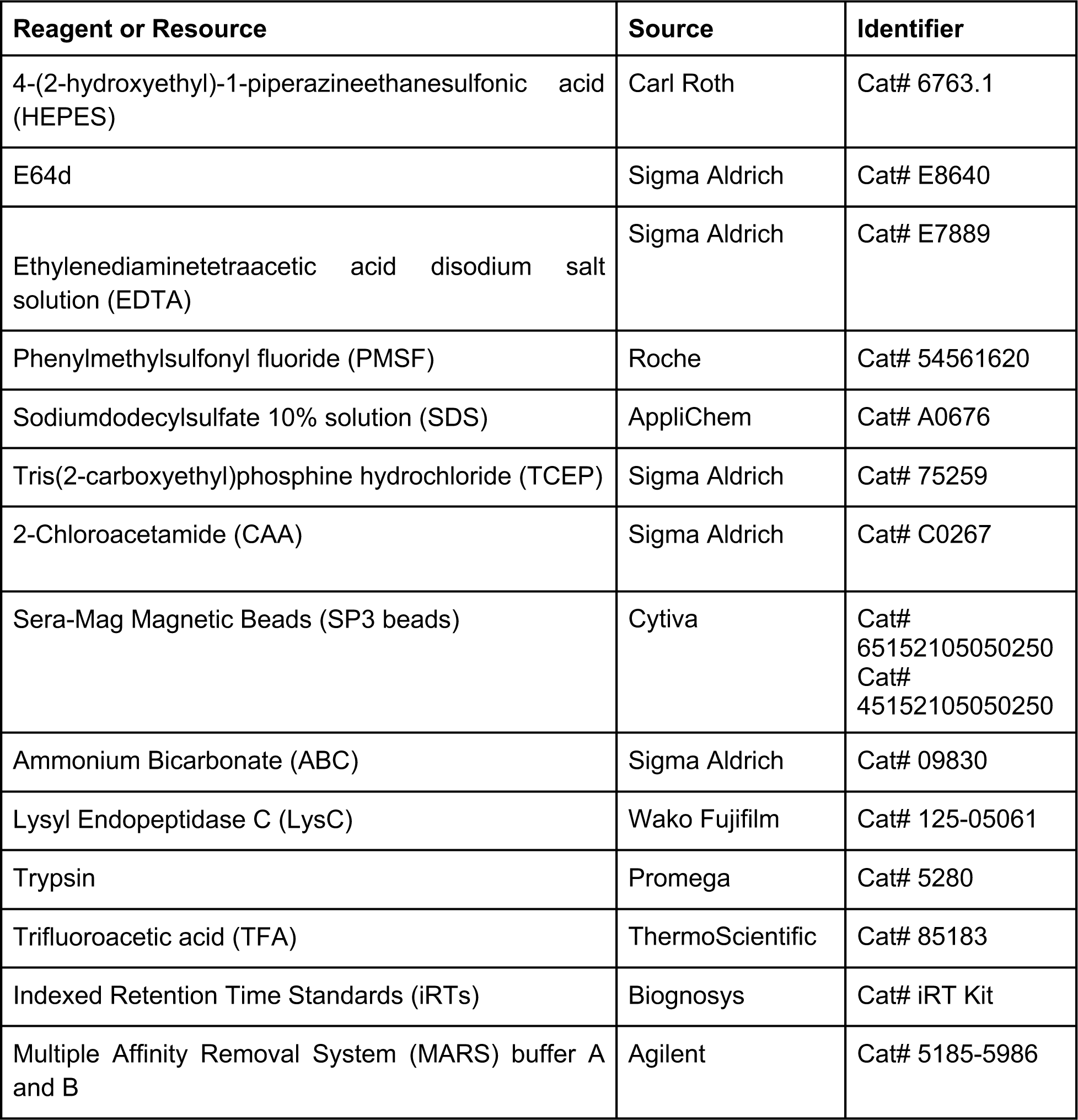
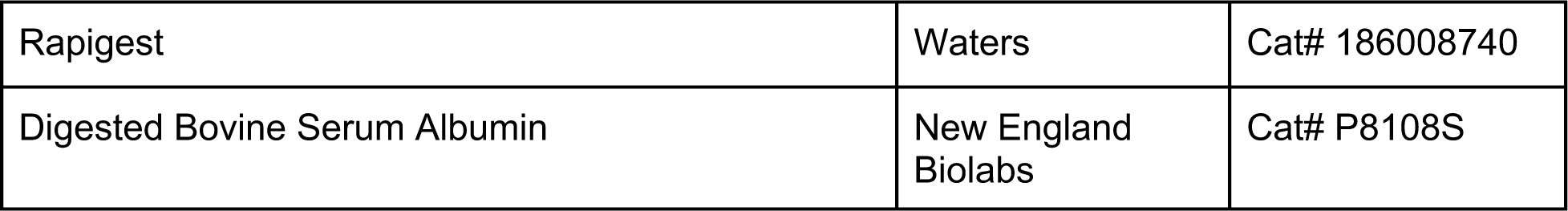

| Reagent or Resource | Source | Identifier |
| --- | --- | --- |
| 4-(2-hydroxyethyl)-1-piperazineethanesulfonic acid (HEPES) | Carl Roth | Cat# 6763.1 |
| E64d | Sigma Aldrich | Cat# E8640 |
| Ethylenediaminetetraacetic acid disodium salt solution (EDTA) | Sigma Aldrich | Cat# E7889 |
| Phenylmethylsulfonyl fluoride (PMSF) | Roche | Cat# 54561620 |
| Sodiumdodecylsulfate 10% solution (SDS) | AppliChem | Cat# A0676 |
| Tris(2-carboxyethyl)phosphine hydrochloride (TCEP) | Sigma Aldrich | Cat# 75259 |
| 2-Chloroacetamide (CAA) | Sigma Aldrich | Cat# C0267 |
| Sera-Mag Magnetic Beads (SP3 beads) | Cytiva | Cat# 65152105050250<br>Cat# 45152105050250 |
| Ammonium Bicarbonate (ABC) | Sigma Aldrich | Cat# 09830 |
| Lysyl Endopeptidase C (LysC) | Wako Fujifilm | Cat# 125-05061 |
| Trypsin | Promega | Cat# 5280 |
| Trifluoroacetic acid (TFA) | ThermoScientific | Cat# 85183 |
| Indexed Retention Time Standards (iRTs) | Biognosys | Cat# iRT Kit |
| Multiple Affinity Removal System (MARS) buffer A and B | Agilent | Cat# 5185-5986 |
| Rapigest | Waters | Cat# 186008740 |
| Digested Bovine Serum Albumin | New England Biolabs | Cat# P8108S |

### Sample Preparation

Using a pair of tweezers, small chunks of frozen tumor tissue were collected in PreCellys tubes (CK14 0.5ml) containing 100ul of 100mM HEPES at pH 8 and a cocktail of protease inhibitors (E64d 1:1000, EDTA 1:50, PMSF 1:100w/v). To prevent thawing, tissue collection was performed on ice. The tweezers were cleaned with 70% Ethanol between each sample. All samples were boiled at 95°C for 10 minutes to deactivate endogenous proteases and then homogenized with a PreCellys 24 homogenizer (4000rpm, 3x30s, 10s pause in between - Bertin Technologies, France). The supernatant was transferred into tubes and remaining beads were washed with 200ul 100mM HEPES pH 8, which was then added to the samples. After addition of SDS to a final concentration of 1% w/v, the samples were sonicated in a Bioruptor Plus for 20 cycles á 30s (Diagenode, Belgium). Resulting protein concentrations were determined with a Bicinchonic acid assay (BCA - Pierce BCA Protein Assay Kit, ThermoFisher, Germany) and 20ug of protein were filled up to 140ul with 100mM HEPES pH 8 and transferred to a 96 deep-well plate. All subsequent steps were performed through an Agilent Bravo liquid handling robot (Agilent Technologies, USA). 5mM TCEP and 20mM CAA were added and incubated for 30 minutes in the dark to ensure reduction and alkylation of proteins. Then 10 ul of a 1:1 mix of washed SP3 beads were added. Acetonitrile was added to reach a final concentration of 50% and mixed on a shaker for 10 minutes at 1050rpm. Afterwards, the beads were separated for 3 min on a magnetic rack and washed twice with 200ul 70% ethanol and once with acetonitrile. During each wash, beads were mixed at 1050rpm for 10min and then separated again on a magnetic rack for 5 minutes. The beads were then taken up in 100ul of 100mM ABC buffer pH 8. Upon addition of 1:100 protein/protease LysC, samples were incubated on a thermoshaker at 47°C for 2h at 600rpm. In a second digestion step, trypsin was added 1:25 protein/protease and incubated overnight at 37°C at 600rpm. The next day, 5% TFA was added to a final concentration of 1%, beads were separated on the magnetic rack and the acidified supernatant collected into a new plate. After peptide concentration determination through a BCA assay, 2ug of peptides were allotted for each sample. 200 fmol of iRTs were added and the samples dried and stored at -80°C.

Frozen GBM serum was thawed, kept on ice and filled up with MARS buffer to a final concentration of 70%. Protease inhibitors EDTA and PMSF were added to final concentrations of 10 mM and 0,1 mM, respectively. After filtration (0,22 um spin filter), samples were depleted semi-automatically with an Agilent Multiple Affinity Removal Column Human 14 column (MARS Cat# 5188-6557, Agilent Technologies, USA) on a microflow HPLC system using MARS buffers A and B in a gradient as suggested by the manual. A BCA was performed to determine the resulting amount of protein. For digestion, Rapigest and HEPES were added to final concentrations of 0,1% and 100mM. Proteins were denatured at 70°C for 10min. Upon addition of final concentrations of 5 mM TCEP and 10 mM CAA, samples were reduced and alkylated for 30 min. 10ul of a 1:1 mix of washed hydrophobic and hydrophilic SP3 beads was added, filled up with acetonitrile to a final concentration of 50% and incubated on a shaker for 10min. The supernatant was removed and beads washed twice in 70% ethanol and once in pure acetonitrile. After removal of remaining acetonitrile, beads were resuspended in 100mM ABC. 1:100 w/w LysC was added relative to previously determined total protein amounts and incubated for 2 hours at 42°C. Then, 1:25 w/w Trypsin was added and incubated overnight at 37°C. Peptide concentrations in the supernatant were determined via BCA. 200 fmol iRTs were added and samples were dried and frozen. From each sample, peptides were also pooled in a mastermix, which was dried and frozen after addition of 200 fmol iRTs per planned injection.

Samples from the meningioma control cohort were prepared and depleted with the same protocol, but digested differently. After depletion and a BCA assay, SDS and HEPES were added to 20 ug of protein to reach final concentrations of 1% and 100mM, and filled up with MARS buffer A. Reduction, Alkylation, SP3-bead binding, washing steps and resuspension of beads in ABC were performed as described for tissues above. Proteins were digested using 1:100 w/w LysC at 42°C for 2 hours and subsequent addition of 1:25 w/w trypsin with incubation at 37°C overnight. Digestion was stopped by adding a final concentration of 1% TFA and drying of the samples. Peptides were resuspended in 0,1% formic acid, 100 fmol of iRTs were added, and 0,5 ug peptides were loaded onto Evotips (Evosep, Denmark) following the Evosep sample loading protocol.

### LC-MS/MS Measurement

GBM tissue samples were analyzed using a LTQ-Orbitrap Fusion Lumos mass spectrometer (Thermo Fisher Scientific, San Jose, CA, USA) coupled to an EASY-nLC 1200 nanoflow liquid chromatography system (Thermo Fisher Scientific (Proxeon), Odense, Denmark). Peptides were loaded directly onto the analytical column and were separated by reversed-phase chromatography using a 50-cm column with an inner diameter of 75 μm, packed with 2 μm C18 particles (Thermo Scientific, San Jose, CA, USA). Chromatographic gradients started at 95% buffer A and 5% buffer B with a flow rate of 300 nl/min for 5 minutes and gradually increased to 25% buffer B and 75% A in 105 min and then to 40% buffer B and 60% A in 15 min. After each analysis, the column was washed for 10 min with 10% buffer A and 90% buffer B. Buffer A contained 0.1% formic acid in water. Buffer B contained 0.1% formic acid in 80% acetonitrile. The mass spectrometer was operated in data-independent acquisition (DIA) mode, with full MS scans over a mass range of 500-900 m/z and detection in the Orbitrap at a resolution of 60,000. The auto gain control (AGC) was set to 2e5 and a maximum injection time of 100ms was used. In each DIA cycle and following each survey scan, 40 consecutive windows of 10 Da each were used to isolate and fragment all precursor ions from 500 to 900 m/z. A normalized collision energy of 30% was used for higher-energy collisional dissociation (HCD) fragmentation. MS2 scan range was set from 350 to 1850 m/z with detection in the Orbitrap at a resolution of 30,000. The AGC was set to 2E5 and a maximum injection time of 60 ms was used. Digested bovine serum albumin was analyzed between each sample to avoid carryover and to assure stability of the instrument. QCloud has been used to control instrument longitudinal performance during the project ^20^.

GBM serum samples were measured on a Thermo QExactive Plus mass spectrometer coupled to a Thermo Easy-nLC 1000 (both ThermoFisher Scientific, USA). Peptides were separated on a uPAC 2m reverse-phase C18 column (ThermoFisher Scientific, USA) using a gradient increasing from 8 to 10 % buffer B in 2 min, then increase to 20 % B within 22 min, to 40 % B in another 46 min, to 55 % B in 10 min, and then to 100 % B in 2 min, where it remains for another 38 min. This 120 min gradient at 300 nl/min was performed with aqueous buffers A containing 0.3 % acetic acid and B containing 80 % acetonitrile and 0,3 % acetic acid. The mastermix was measured in six gas-phase fractionation measurements, which covered 100 m/z ranges between 400 and 1000 m/z in 25 staggered acquisition windows of 4 m/z. Sample measurements were performed in a range from 385 to 1015 m/z in staggered 24 m/z acquisition windows. We used a resolution of 17,500, 80 ms maximum injection time and Higher-energy C-trap dissociation (HCD) in stepped normalized collision energies of 25 and 30.

Meningioma control samples were measured on a Bruker TimsTOF Flex (Bruker, USA) connected to an Evosep One HPLC system (Evosep, Denmark). Samples loaded on Evotips were separated with the vendor’s standard 30 samples per day (30 SPD) method using a 44min gradient at a flow of 500 nl/min and an EV 1137 performance 15cm reverse-phase column. The mass-spectrometer was equipped with a Captive Spray ion source and operated at a mass range from 400 to 1200 m/z and a mobility range from 0.6 to 1.6 1/K0. DIA measurements were performed in 32 mass windows of 26 m/z with an overlap of 1 m/z. The cycle time was set at 1.8s and collision energies ranged from 20 to 59eV.

### Data Analysis

Peptide to spectrum matching of GBM tissue MS raw data was performed in DIA-NN 1.8, and for GBM serum samples DIA-NN 1.7.12 was used ^21^. For GBM tissue samples, a library was predicted using a human reference proteome as downloaded from uniprot.org on 03.03.2022, refined on all samples of the cohort, and filtered at 1% false discovery rate (FDR). Only peptides between 7 to 30 amino acids length and within 500 and 900 m/z were included, but one missed cleavage site was allowed. Samples were reannotated using a 1% FDR cutoff. For GBM serum samples, a predicted library using the same reference proteome as above was refined against the gas-phase fractionation measurements of the mastermix and filtered at 10% FDR. Peptide lengths from 7 to 30 amino acids were allowed and precursor masses ranging from 400 to 1000 m/z considered. In both DIA-NN analyses, up to one missed cleavage site were considered and Reannotation was performed at 1% FDR. Raw data from the meningioma control cohort was analyzed in Spectronaut (Biognosys, Switzerland) with the same human reference proteome using the software’s custom directDIA workflow ^22^. Also here, a FDR threshold of 1% was used and only peptides ranging from 7 to 52 amino acids within a mass range from 400 to 1200 m/z considered. Here, we allowed for up to two missed cleavage sites.

All subsequent statistical analysis was performed in R version 4.2 and R Studio. A protein expression matrix was generated from the DIA-NN output by using the DIA-NN R package ^21^. Here, quantification of peptides and inferred proteins was performed through the MaxLFQ algorithm by only considering proteotypic peptides ^23^. All protein intensities were then median centered and log2 transformed. For the semi-specific analysis, we used the same DIA-NN workflow, but with a custom-made reference proteome already containing semi-tryptic peptide sequences.

Subsequent statistical analyses were conducted using in-house scripts and publicly available R packages. Unsupervised statistical analyses were done with the MixOmics ^24^. The optimal number of hierarchical clusters was determined with a Monte-Carlo simulation using M3C ^25^. For differential expression analyses we used the Limma package and clusterProfiler for gene set enrichment and overrepresentation analyses together with the KEGG, Reactome, and Gene Ontology databases ^26–32^. Imputations were performed with MissForest or DIMA ^33,34^. Semi-specific peptides were annotated with the Fragterminomics package ^35^.

## Results and Discussion

### Cohort Overview

Our cohort consisted of matching tissue and serum samples from 55 GBM patients. The median age was 66 years (interquartile range IQR = 61 - 75) and 42% of the patients were female (Table 1). Included were primary tumor tissues from all patients and in five cases samples from matching recurrent tumors, depending on the time to recurrence. Serum was collected one day before (preOP) and three days after (postOP) surgery. Additionally, we assembled an independent cohort of serum samples from 26 Meningioma (MNG) patients collected in the same way (Fig. 1A, B). These samples were used to control for bias introduced by brain surgery.

**Fig. 1.**
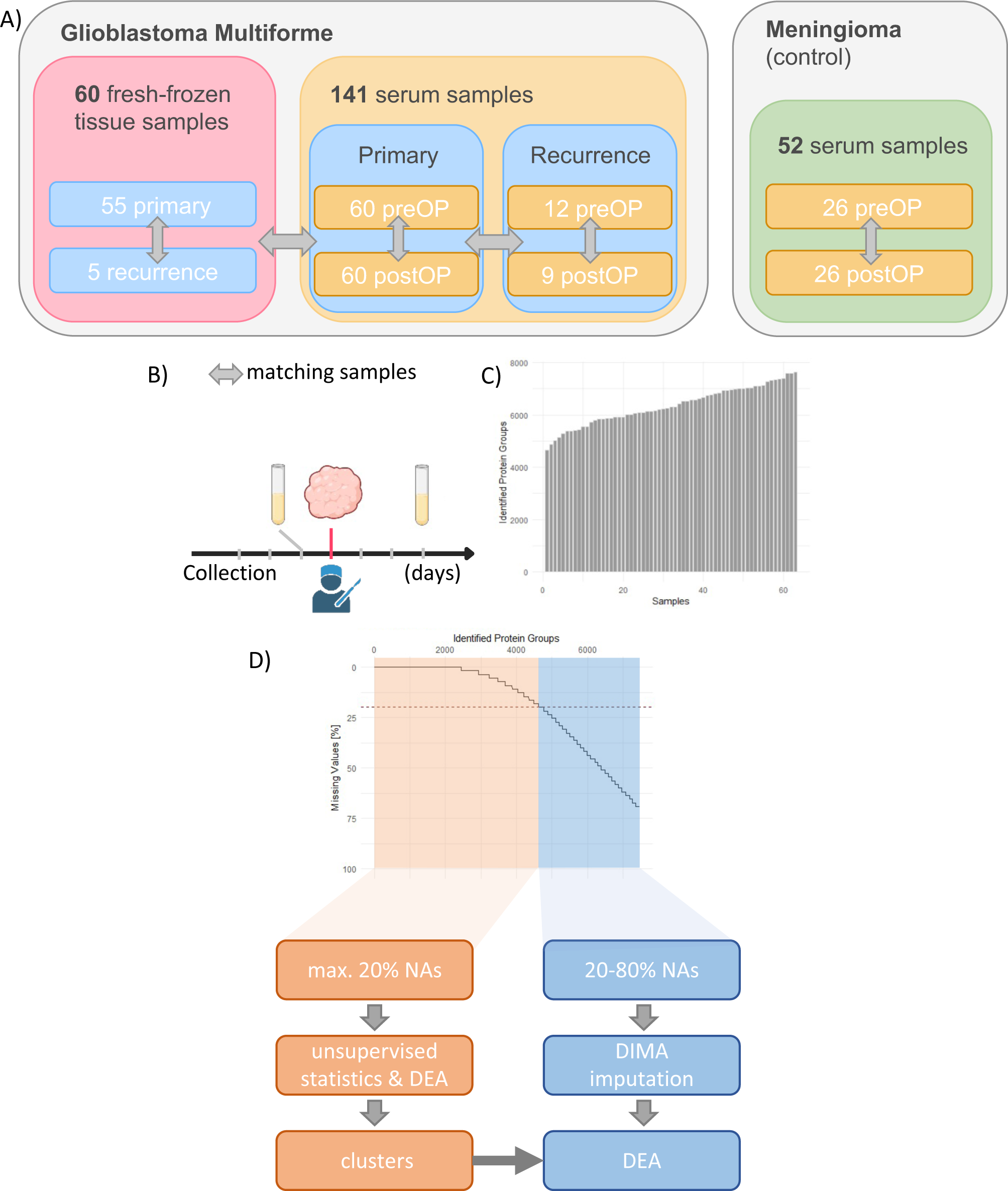
GBM cohort. A) Setup of the matching GBM tissue and serum cohorts together with independent Meningeoma control cohort. B) Time-points for sample collection. C) Identified proteins in GBM tissue cohort. D) Missingness in GBM tissue cohort and resulting analysis strategies for proteins with low and high missingess. NA - missing value. DEA - Differential Expression Analysis.

**Tables.**
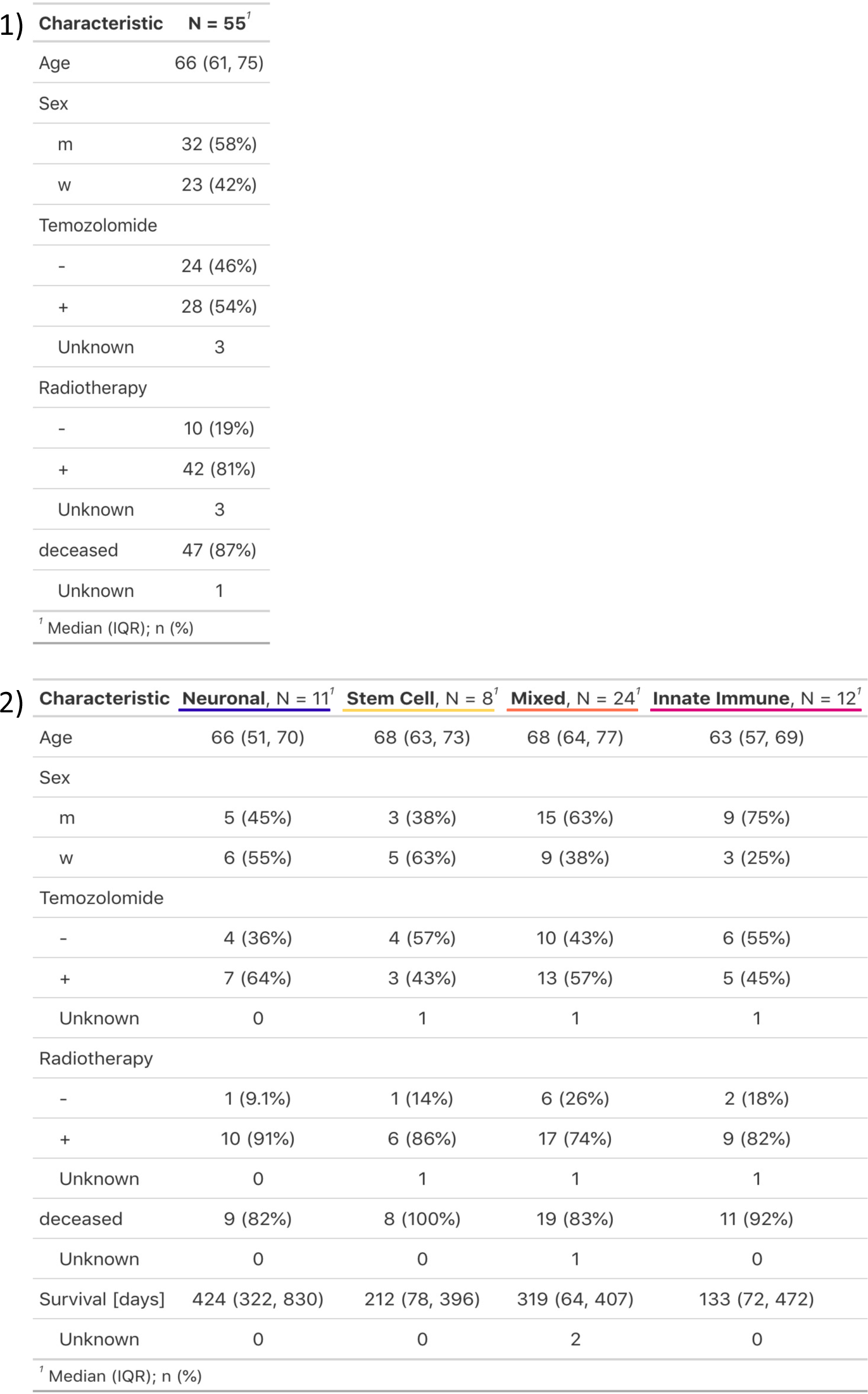
1) Clinical annotation of cohort. 2) Clinical annotation per cluster. IQR – interquartile range.

|  |  |
| --- | --- |
| Age | 66 (61, 75) |
| Sex |  |
| m | 32 (58%) |
| w | 23 (42%) |
| Temozolomide |  |
| - | 24 (46%) |
| + | 28 (54%) |
| Unknown | 3 |
| Radiotherapy |  |
| - | 10 (19%) |
| + | 42 (81%) |
| Unknown | 3 |
| deceased | 47 (87%) |
| Unknown | 1 |

<sup>†</sup> Median (IQR); n (%)

|  |  |  |  |  |
| --- | --- | --- | --- | --- |
| Age | 66 (51, 70) | 68 (63, 73) | 68 (64, 77) | 63 (57, 69) |
| Sex |  |  |  |  |
| m | 5 (45%) | 3 (38%) | 15 (63%) | 9 (75%) |
| w | 6 (55%) | 5 (63%) | 9 (38%) | 3 (25%) |
| Temozolomide |  |  |  |  |
| - | 4 (36%) | 4 (57%) | 10 (43%) | 6 (55%) |
| + | 7 (64%) | 3 (43%) | 13 (57%) | 5 (45%) |
| Unknown | 0 | 1 | 1 | 1 |
| Radiotherapy |  |  |  |  |
| - | 1 (9.1%) | 1 (14%) | 6 (26%) | 2 (18%) |
| + | 10 (91%) | 6 (86%) | 17 (74%) | 9 (82%) |
| Unknown | 0 | 1 | 1 | 1 |
| deceased | 9 (82%) | 8 (100%) | 19 (83%) | 11 (92%) |
| Unknown | 0 | 0 | 1 | 0 |
| Survival [days] | 424 (322, 830) | 212 (78, 396) | 319 (64, 407) | 133 (72, 472) |
| Unknown | 0 | 0 | 2 | 0 |

<sup>†</sup> Median (IQR); n (%)

### High proteome coverage in GBM tissue

We identified and quantified more than 6300 protein groups on average per GBM tissue sample (Fig. 1C). More than 4600 of these proteins were present in 80% of all samples and used for initial statistical analysis. 2500 additional proteins were detected in less than 80%, but more than 30%, of samples and used for a missingness analysis as described below (Fig. 1D). Batch effects from sample preparation or measurement order were not apparent (Fig. S1).

### Immune infiltration and neuronal development markers define four proteomic GBM clusters

At first, we considered 55 primary tumors alone and used hierarchical clustering to define proteomic subtypes of primary GBM (Fig. S2A). A Monte-Carlo simulation did not fortify a specific number of optimal subclusters (Fig. S2B). We decided to split the data into 4 subtypes with meaningful biological motifs: a neuronal cluster, an innate immunity cluster, a mixed cluster, and a stem cell cluster (Fig. 2A). These subtypes did not correspond to *EGFR* variant vIII and p53 accumulation, *ATRX* mutation or Ki67 levels as previously determined by neuropathological routine analyses (Fig. S2A). In a principal component analysis (PCA) and uniform manifold approximation and projection (UMAP), we noticed a distinct separation of the neuronal cluster from the other clusters. Conversely, the innate immunity, mixed, and stem-cell clusters exhibited overlapping features and transitioned into each other (Fig. 2C & 6A). A differential expression analysis comparing each cluster to all other samples, recapitulated their gradual and interconnected nature (Fig. S2B). Because cancer biology is shaped by interactions of the tumor with the immune system and GBM might arise from malignant transformation of neuronal stem cells, we also screened our data for immune- and neurodevelopmental markers ^36,37^.

**Fig. 2.**
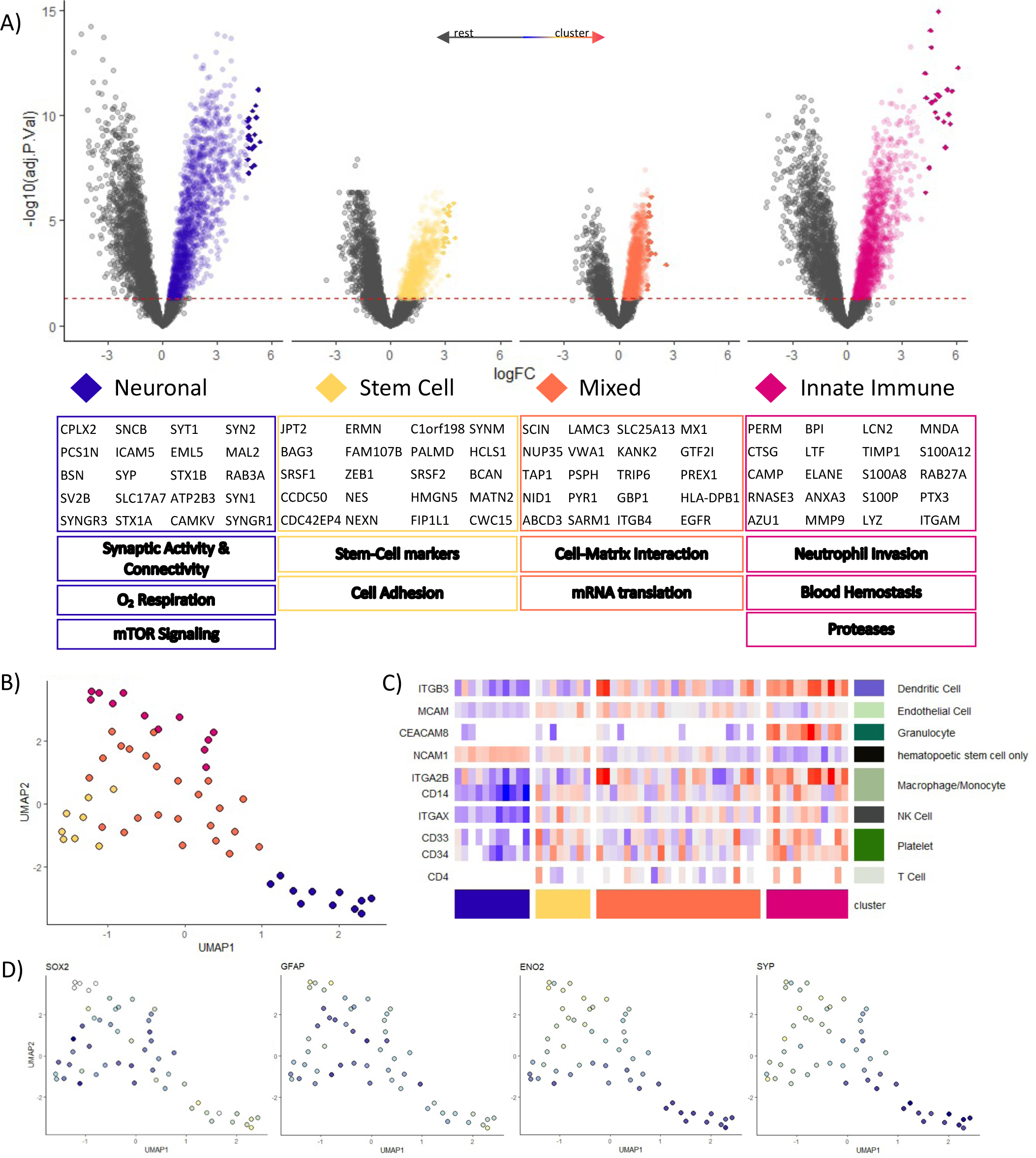
Unsupervised statistics and clustering of primary tumor tissues. A) Volcano plot of differential expression analysis one cluster vs. all other clusters (rest). The 20 most upregulated proteins per cluster are shown in diamonds. Gene names of these proteins and globally enriched cellular processes are given below. B) UMAP of clusters C) Heatmap of specific immune cell markers. D) UMAP expression profiles of selected neurogenesis markers

To some extent, the clusters reflect expression patterns that were described in previous studies ^11,14–18^. The neuronal cluster’s high abundance of proteins linked to synaptic activity, connectivity, and increased aerobic respiration matches well to a proneural-like subtype. Besides a noteworthy upregulation of NTRK2 and -3, it also showed an mTOR signaling fingerprint, including the oncoproteins HRAS and KRAS. Tumor suppressor PTEN, which can be deleted during chromosomal rearrangement, was expressed at higher levels than in all other clusters ^38^. The cluster consistently presented the lowest abundance levels for immune markers. Only proteins like NCAM1 or L1CAM, which are also canonically expressed in the brain, were enriched here (data from v23.proteinatlas.org). Conversely, we found that MAP2 and ENO2, as well as synapse markers SYP and SYT1, which all signify mature and functional neurons, were only upregulated in the neuronal cluster (Fig. 2D & S4A) ^39–42^.

The mixed cluster, which included the most patients, cannot be clearly demarcated from the innate immune and stem cell clusters and might represent an intermediate state. Due to its higher expression of EGFR and mRNA translation and cell-matrix interaction related proteins, it resembles a classical subtype ^14^. It only showed a moderate immune cell infiltration. Proteins like GFAP, SOX10 or BLBP, which indicate differentiation towards glial cells, were most present in this cluster (Fig. 2D & S4A) ^43,44^.

Similarly, the stem cell cluster shows a high mRNA splicing and processing activity, but also an upregulation of stem cell markers. The neuronal progenitor cell marker NES and transcriptional repressor ZEB1 are highly expressed here, as well as early stem cell markers SOX2 and NOTCH1 ^45,46^. Tumor suppressor NF1 was downregulated ^14,38^. ZEB1 and SOX2 have been linked to increased glioblastoma invasion and therapy resistance ^47–49^. Immune markers were detected here as well, but only at intermediate levels (Fig. 2D & S4A).

Lastly, the innate immune cluster showed a pronounced upregulation of immunity and hemostasis related proteins, especially of neutrophil proteases like CTSG and LYZ (Fig. 2A). Unsurprisingly, from a list of specific immune markers, 8 out of 10 were detected predominantly here (Fig. 2C). Especially granulocyte-, macrophage / monocyte-, and dendritic cell markers were increased and we also found a high expression of MRC1 (CD206) and CD163, which both indicate the presence of M2 polarized macrophages ^50^. A more expansive list of less cell-type specific immune markers confirmed this pattern (Fig. S3).

Overall, our data suggest that the GBM proteome is molded by the combined impact of neuronal differentiation and tumor immune reactions. Further investigations are necessary to clarify whether our data reflects regional protein expression differences within the tumors or actually represents mutually exclusive proteomic subtypes.

### Missingness analysis confirms and expands cluster’s biological motifs

Since feature reannotation in mass spectrometry-based proteomics is dependent on the signal intensity, non-detection of a peptide / protein does not automatically mean its absence ^51,52^. Because DIA measurements ensure good identification consistency across cohorts, missing values at least imply expression levels below the detection limit if the peptide was found in other samples ^51^. Thus, the distribution of missing values can contain valuable information, although it is not directly accessible to classical statistical approaches. To investigate whether proteins with higher missingness were detected selectively only in some clusters, we imputed missing values with the DIMA algorithm and extracted proteins which had shown no more than 70% missingness before imputation ^34^. We then tested for differentially abundant proteins between the clusters with a focus on proteins that had not been discerned in the initial approach (Fig. 1D). We found 2533 proteins that showed significant differences in abundance and missing value distribution between the clusters. Overrepresentation analyses confirmed and expanded already defined biological motifs for the clusters. For example, mTOR signaling components emerged to be highly enriched in the mixed cluster. This approach also enabled us to allocate disease relevant proteins with high missingness into the clusters, as was the case for pattern-recognition receptor Ficolin 1 (FCN1), which was more abundant in the innate immunity cluster, or the neuronal cluster enrichment of gamma-aminobutyric acid (GABA) receptor 2 GABRG2. In the same way, cell proliferation marker Ki67 was detected well in the mixed-, innate immune-, and stem cell cluster, but not at all in the neuronal cluster (Fig. 3A&B). This is in contrast to results obtained via immunohistochemistry, which display a rather uniform Ki67 expression (Fig. 3C).

**Fig. 3.**
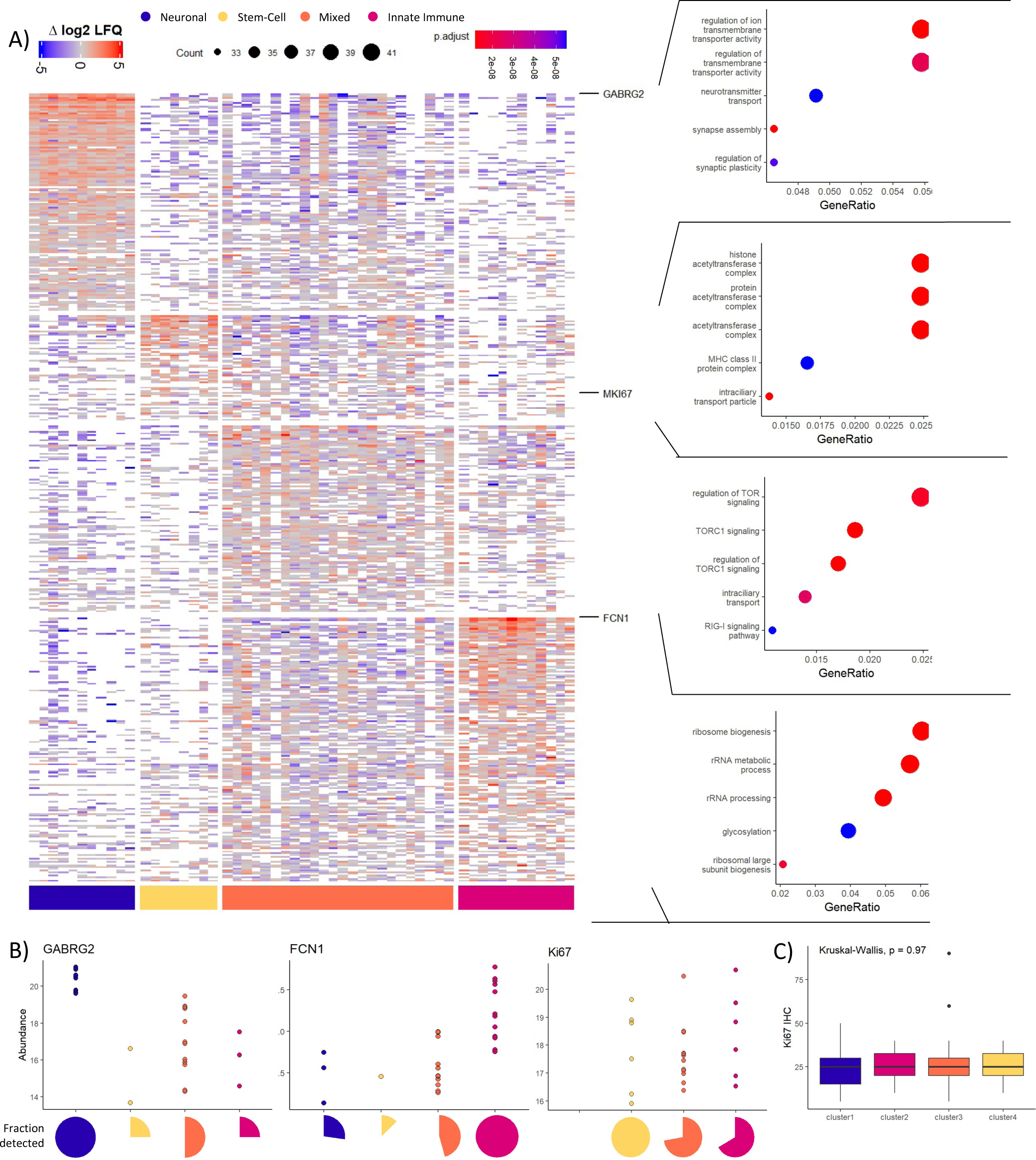
Missingness Analysis. A) Left: Heatmap of proteins with 20-70% NAs as discerned in missingness analysis. NA values are left in white. Right: Overrepresentation analysis results per group B) LFQ abundance and coverage per cluster for selected proteins. C) Ki67 expression levels as determined via immunohistochemistry.

### Overall survival significantly longer in neuronal cluster than in stem cell cluster

Prolonging patient survival is the crucial aspect of GBM therapy. In our cohort, patients had a median survival of 336 days and 87% of patients died within the study duration of 1624 days (Tables 1 & 2, Fig. 4A). Survival probability was dependent on a range of different factors. In agreement with previously published data, elderly patients have a significantly increased risk for earlier death (HR 1.04, p = 0.008, Fig. 4B) ^3,53,54^. The patient’s gender did not have a significant impact on survival (log rank p = 0.47, Fig. 4E). Likewise, a complete resection of the primary tumor might give a short survival advantage after surgery, but did not change the probability of survival throughout the whole duration of the study (log rank p = 0.5, Fig. 4F). The choice of therapy, however, proved to be decisive, as patients who opted for radio- and/or chemotherapy with TMZ at some point in their treatment, survived significantly longer (log rank radiotherapy p = 0.0063; TMZ p = 0.00012, Fig. 4C & D) ^53,54^. Notably, the tumor’s assigned cluster had an impact as well. While patients of the stem cell-, mixed-, and innate immune cluster all showed similar survival, patients of the neuronal cluster tended to survive longer (log rank p = 0.09, Fig. 4G). Similarly, in Cox proportional hazards model (CPHM), the stem cell cluster shows significantly worse prognosis relative to the neuronal cluster (HR 3.4, p = 0.017, Fig. 4H). A potential explanation for these differences in survival are the lower degree of inflammation and lower expression of neurogenesis markers in the neuronal cluster. A survival analysis on protein level with Age, TMZ- and radiotherapy as mandatory covariables via the CoxBoost R package or multiple-testing corrected CPHM did not reveal patterns of individual proteins to correlate with survival (Fig. S4B) ^55^.

**Fig. 4.**
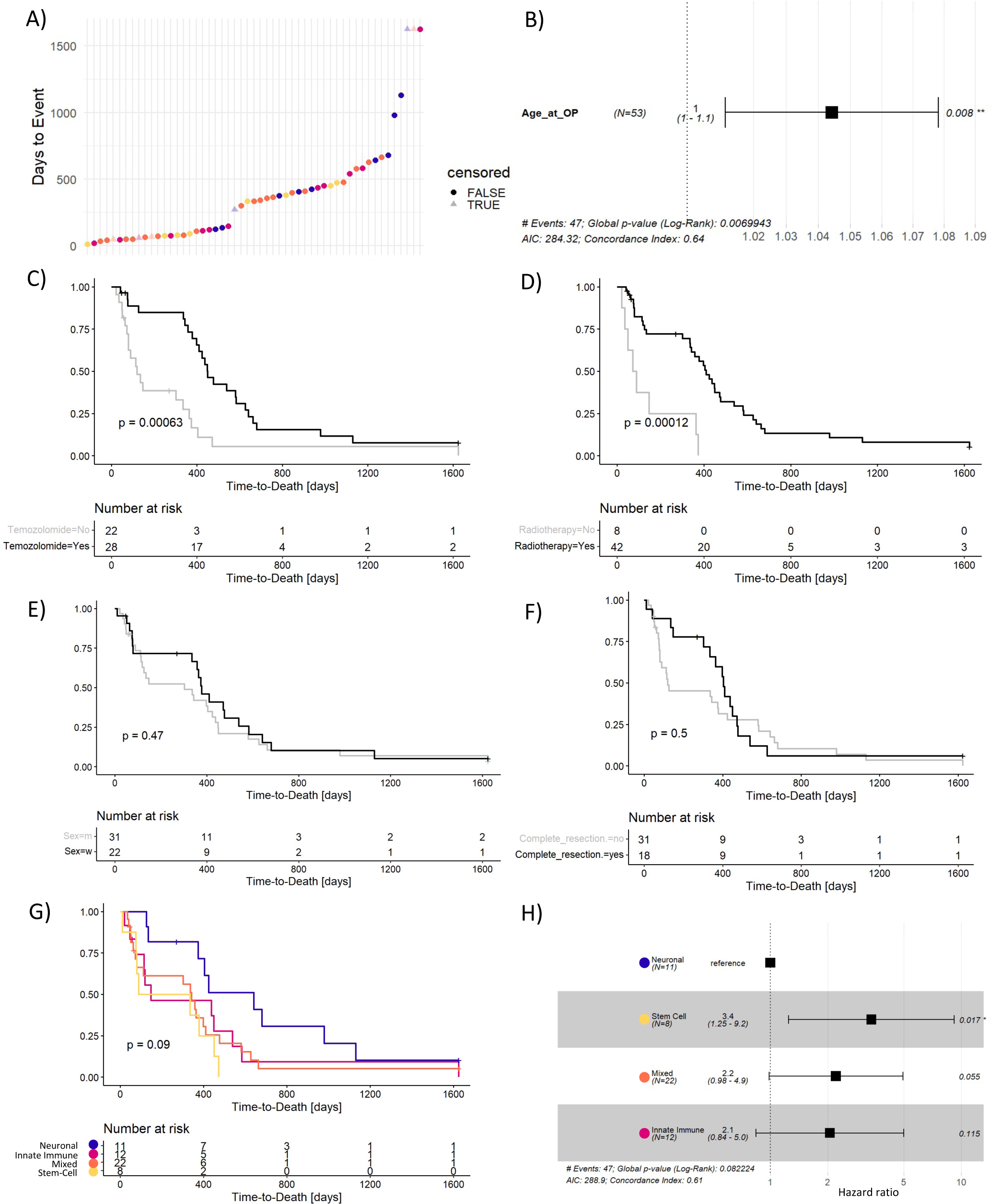
Survival Statistics. A) Distribution of overall survival throughout cohort. Censored values are shown as triangles. B) Forest plot of hazard ratio for Age at primary surgery. C - G) Kaplan-Meier plots including log rank test significance for TMZ treatment, radiotherapy, sex, complete resection, and assigned cluster. H) Forest plot of hazard ratios for different clusters.

### Elevated proteolytic activity in the innate immune cluster

In a second, semi-specific analysis of our data, we searched for peptides that underwent proteolytic cleavage within the tumor and thus only possessed one tryptic terminus. In total, we identified 19876 of such semi-specific proteolytic products, which represented 22% of all detected peptides (Fig. 5A). The fraction of semi-specific peptides was significantly increased in tumors from the innate immunity cluster, which fits to the high expression of (neutrophil) proteases in these tissues (Fig. 5B). When comparing abundances of the full protein with corresponding proteolytic products in each cluster, most proteins appear to be cleaved relative to their overall expression level supposedly due to unspecific protein degradation in a highly proteolytic environment. Some semi-specific peptides, however, were disproportionately high- or low abundant compared to their protein of origin, which suggests regulated proteolytic processing (Fig. 5C). One physiologically relevant proteolytic product with a size and cleavage site similar to the peptide observed in our data, was e.g. already described for RTN4 ^56–58^.

**Fig. 5.**
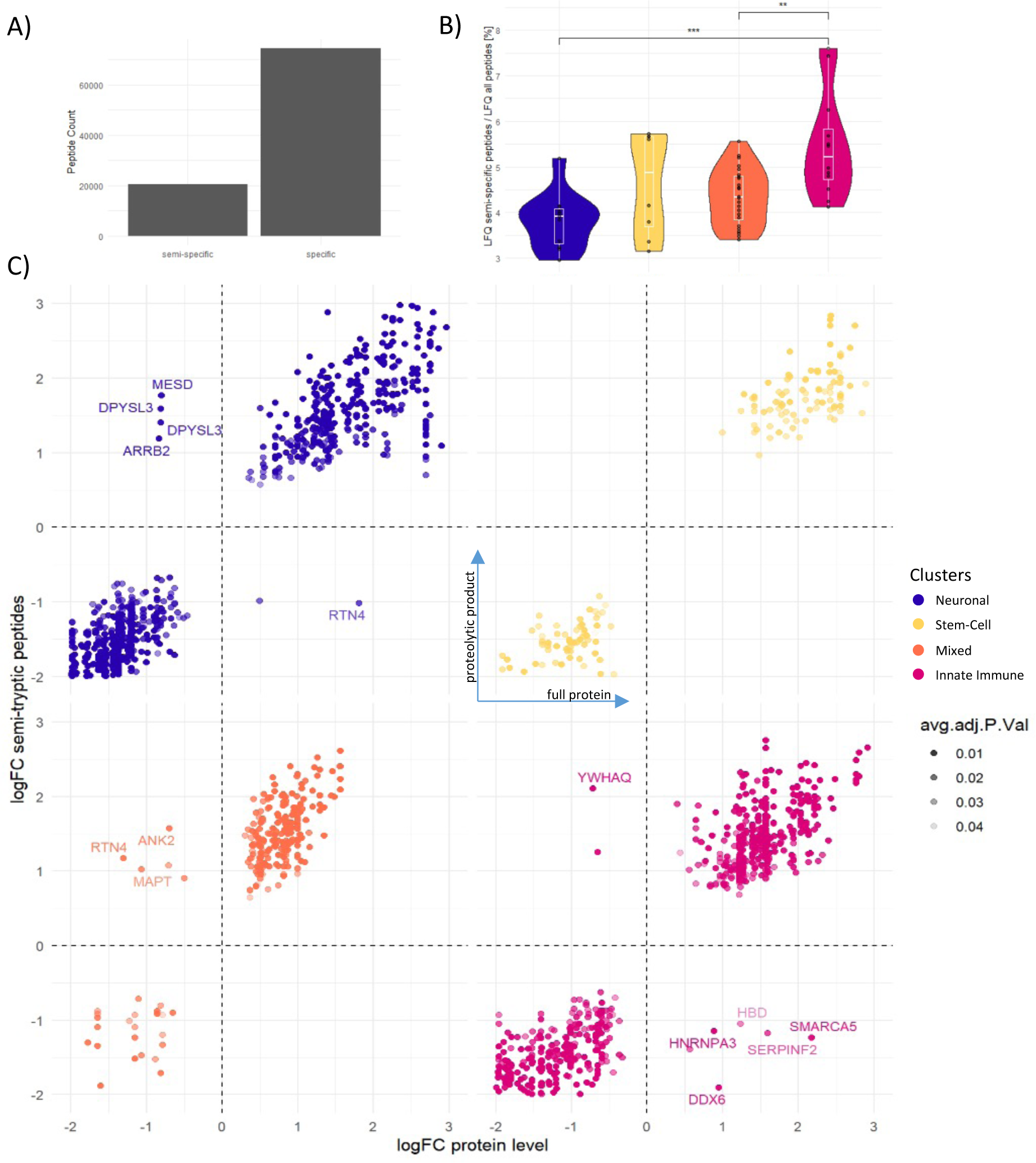
Semi-specific analysis of primary tumor tissues. A) Identified semi-specific and fully tryptic peptides. B) Fraction of detected semi-specific peptides per cluster. C) Correlation of fold-changes between protein abundance and corresponding semi-specific peptides.

### Matching recurrences show distinct protein expression profiles

As previously mentioned, our cohort contained five matching pairs of primary and recurrent tumors. Although recurrences did not cluster separately from all primary tumors in hierarchical clustering or PCA, they showed systematic protein expression changes when compared to their matching primary tissues (Fig. 6A). A partial least-squares discriminant analysis (PLS-DA) further highlighted systematic proteome differences (Fig. 6B & S5A). In a patient-matched, differential expression analysis, proteins involved in innate immunity and inflammation were downregulated in recurrences, while neuronal growth regulation and synaptic activity was upregulated (Fig. 6D, E). These findings point towards proteomic plasticity of GBM recurrence. Buehler et al. also report dampened inflammation in recurrent GBM while Cosenza-Contreras et al. observed the opposite trend ^59,60^. A semi-specific analysis of the tissues revealed no difference in proteolytic activity between matching primary-recurrence pairs (Fig. S5B). However, in differential expression analyses comparing recurrent vs. primary tumors, we observed that fold changes of some semi-specific peptides did not match the abundance of their protein of origin, which possibly indicates altered proteolytic processing (Fig. 6C).

**Fig. 6.**
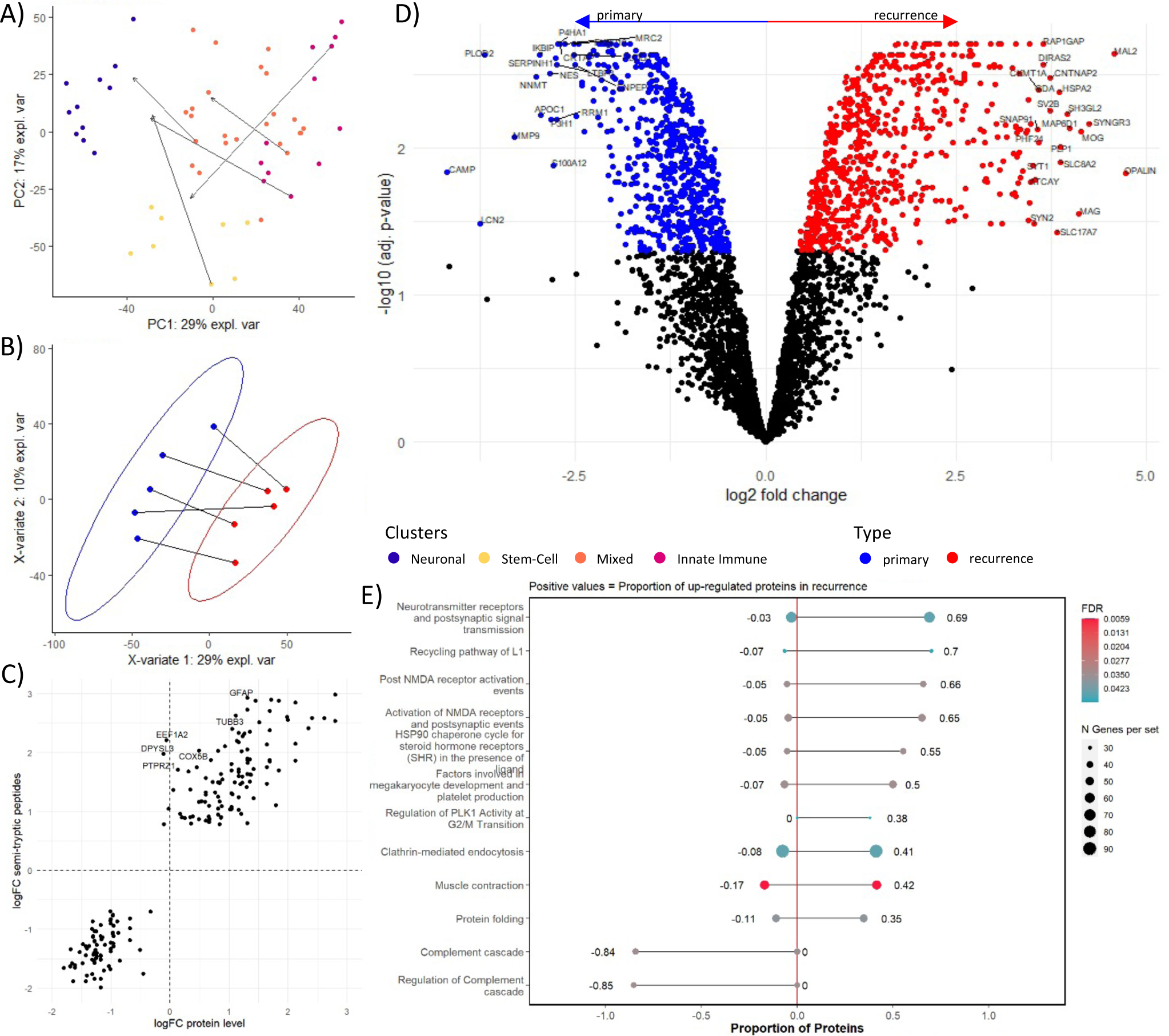
Recurrent vs. primary tumor tissues. Primary tumors are shown in blue, recurrent tumors in red. A) PCA of the entire GBM tissue cohort with colors indicating the primary tumor clusters. Arrows show placement of matching recurrences. B) PLS-DA of matching primary-recurrent tissue pairs. C) Correlation of fold-changes between protein abundance and corresponding semi-specific peptides for matching primary vs. recurrent samples. D) Volcano plot of patient-matched differential expression analysis recurrent vs. primary tumors. E) Gene Set Enrichment Analysis showing biological processes upregulated in primary and recurrent tumors.

### Overview of GBM serum profiling

In the GBM serum cohort, we identified on average 735 proteins after semi-automated depletion of 14 highly abundant serum proteins (Fig. 7A). 549 of these were present with less than 20% missingness (Fig. 7B). To control for biases introduced by surgery, we also included an independent cohort of preOP and postOP serum samples from 26 Meningioma (MNG) patients. These samples had been collected using the same time points. MNG are slow-growing brain tumors. Although they do not share GBM’s highly proliferative, invasive, and heterogeneous nature, patients still undergo brain surgery with similar protocols ^61,62^. Slightly higher protein identification numbers were recorded in the MNG control cohort, which was depleted with the same workflow, but digested and measured using different protocols and devices (Fig. S6A-C). MNG serum samples presented very similar to GBM serum in their individuality, the systematic upregulation of mostly the same inflammatory markers, and overall proteome changes post-surgery. This similarity also became evident, when we merged both cohorts via the ComBat algorithm, which led to a nearly perfect overlap (Fig. S6F) ^63^.

**Fig. 7.**
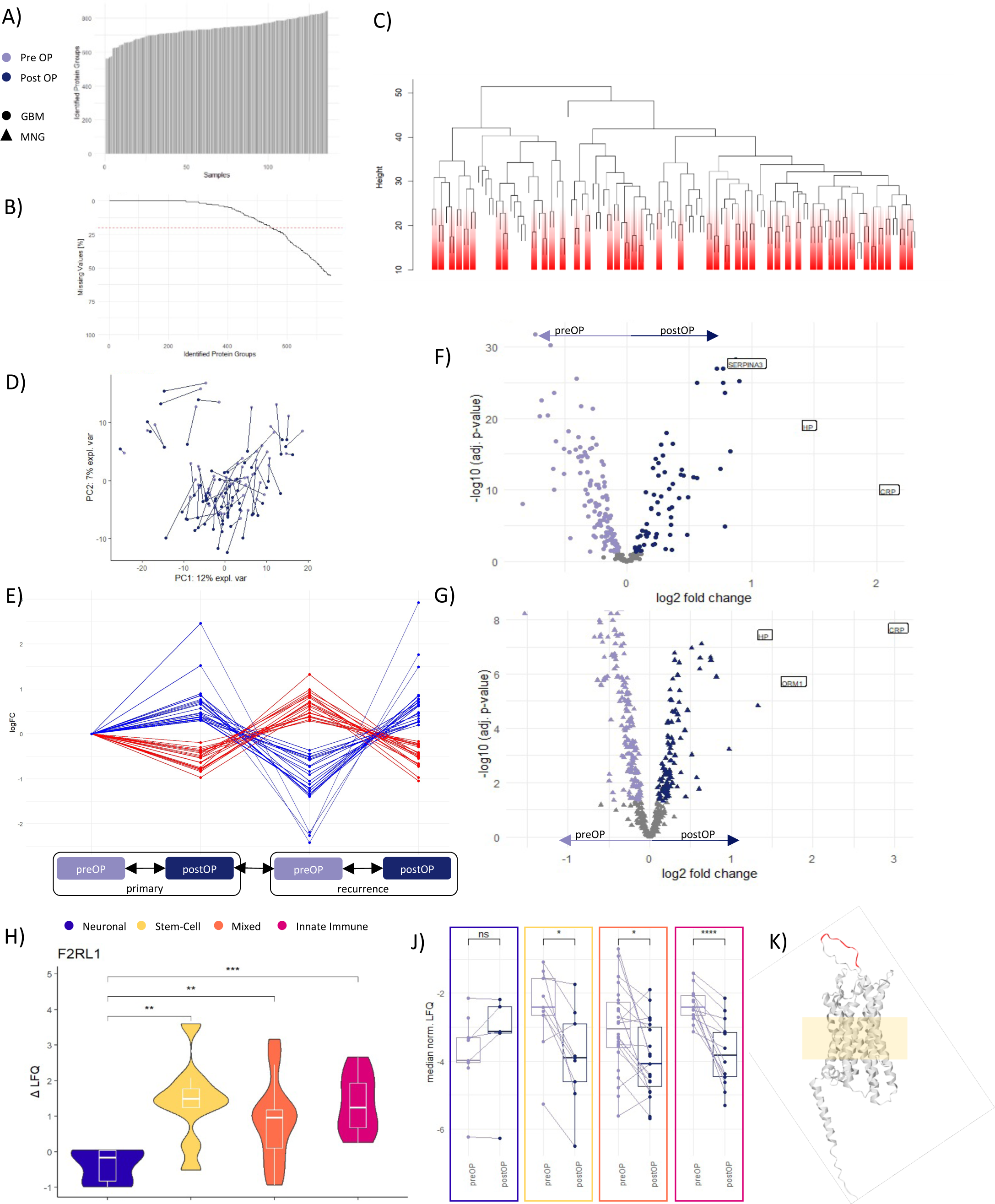
Patient Serum. A) Identified proteins in GBM serum cohort. B) Missingness in GBM serum cohort. C) Hierarchical clustering of GBM serum cohort. Pre- and post OP samples from the same patient clustering together are given in red. D) PCA of GBM serum cohort. Matching samples are connected. E) Patient-matched differential expression analysis results comparing matching preOP and postOP serum from patients with primary and recurrent tumors. F & G) Volcano plots of patient-matched differential expression analysis post OP vs. pre OP (top: GBM serum cohort. bottom: Meningioma serum cohort). Acute phase proteins are labelled. H) ΔLogFC between matching serum samples from primary and recurrent tumors. J) Change of F2RL1/PAR2 abundance in GBM serum cohort between clusters. Matching sample pairs are connected. K)

### Serum proteome profiles are highly individual

In PCA and hierarchical clustering of GBM serum, samples from the same patients frequently clustered together, despite narcosis, surgery, and recovery in between collection points (Fig. 7C,D). This reflects earlier results by Liu et al. and highlights the remarkable individuality and stability of the human serum proteome ^64^. After surgery, mostly inflammatory processes and particularly acute phase proteins were upregulated in a patient-matched differential expression analysis, which was mirrored by the MNG control cohort (Fig. 7F,G).

### The quest for a GBM-specific serum protein signature

One main objective of our work was to find circulating markers that are associated with GBM recurrence. To select candidate proteins, we performed a multi-tiered approach with the following criteria: I) an expression pattern defined as a drop in abundance after tumor removal and increased abundance prior to recurrence surgery, II) published overexpression in GBM tissue relative to non-malignant brain. We also compared expression changes to protein levels in MNG serum to test for GBM specificity and control for confounding effects of the surgery. Additionally, we searched for proteins with correlating expression levels in tumor tissue and serum. We employed different data analysis strategies to identify proteins with such properties.

### Marker Candidates with specific Expression Pattern

At first we investigated for proteins, which decreased in abundance after the tumors were removed, but increased in between surgeries. Matching preOP and postOP serum samples from primary and recurrent tumors were available from 9 patients, which we compared in multiple patient-matched differential expression analyses. In a second step, we controlled that selected proteins also decreased after primary surgery in the entire cohort of 55 patients. We identified 20 proteins, which followed this expression pattern, and 21 proteins, which showed inverse expression. 27 of these 41 proteins behave similarly in the MNG cohort and are thus not GBM specific. However, they might provide perspectives for larger GBM serum studies of circulating markers for recurrence, if controlled for confounding effects of brain surgery. 10 more were only detected in GBM, but not MNG samples, despite higher peptide and protein coverage in the MNG cohort. Three of the proteins exhibited differing expression patterns between GBM and MNG cohorts (all Fig. 7E & Table S1). PLA2G7 and, to a lesser extent, LUM did not significantly change expression in the MNG cohort, but followed the specified expression pattern in GBM. The phospholipase PLA2G7 is an established marker for cardiac diseases and was proposed as a circulating and tissue-resident marker for a variety of cancers ^65–69^. The protein is predominantly expressed by leukocytes (data from v23.proteinatlas.org) and was detected in only 6 GBM tissue samples. LUM overexpression in GBM tissue was already reported on the transcriptomic level (Fig. S7A) and in a proteomic dataset from our group ^70^. Conversely, ALB strictly followed an inverse expression pattern, meaning it increased significantly after surgery in the GBM, but not in the MNG cohort. Albumin, which normally makes up more than 50% of the serum proteome, was depleted from the samples, but still remained detectable. Serum albumin levels respond to many physiological stimuli, including inflammation and surgical trauma ^71^. Serum albumin is also a known prognostic marker for GBM and might thus be regulated differently between both cohorts^71–74^.

### Apolipoprotein(a) *levels correlate between tissue and serum*

We then searched for proteins whose abundance in serum correlated to their tissue expression levels. 299 proteins were sufficiently detected with less than 20% missingness in GBM serum and tissue. Remaining missing values were imputed with the MissForest algorithm. When comparing preOP or postOP serum to tissue, all except one protein had weak Pearson correlation coefficients below 0.5 (Fig. S6E,F). Only cardiovascular risk marker Apolipoprotein(a) (LPA) correlated well in both comparisons (correlation coefficient preOP-tissue = 0.76; postOP-tissue 0.72). Since LPA expression rates are considered to be predominantly determined by genetic inheritance, we presume that this reflects documented inter-individual expression variations exceeding a thousand-fold rather than tumor-intrinsic properties ^75,76^.

### PAR2 abundance in highly individual serum proteomes matches to tissue clusters

To account for individual differences in serum proteome compositions, proteome alterations due to surgery, and molecular subtypes of the tumor tissue, we next calculated the spread between postOP and preOP protein abundances (ΔLFQ) and compared these values between the matching tissue clusters. Here, we found one peptide from proteolysis activated receptor 2 (PAR2/ *F2RL1*) to consistently drop after surgery in innate immune- and, to a lesser extent, stem-cell- and mixed cluster patients (Fig. 7H,J & S6F). PAR2 is highly expressed in neutrophils and the brain, where it directs a crucial pathway for the induction of neuroinflammation ^77,78^. *F2RL1* mRNA is also overexpressed in GBM tissue (Fig. S7B). PAR2 senses proteolytic environments through its extracellular N-terminus, which auto-activates the receptor when cleaved ^78,79^. The detected circulating peptide is part of this N-terminus and its abundance reflects matching tumor tissue’s proteolytic activity. Removing the tumor might eliminate PAR2-expressing tissue and secreted proteases, making circulating PAR2-peptide levels a potential indicator for the inflamed tumor mass. The levels of the circulating PAR2-peptide were very individual, but followed the same trajectory. Closely monitoring PAR2-peptide levels in a patient before and after surgery might thus allow the detection of regrowing tumors, if they become inflamed. PAR2 expression is not specific to GBM and fragments have been detected in other diseases ^80^. Despite a much higher peptide coverage, we did not identify any circulating PAR2 peptides in the Meningioma control cohort. Neither did we detect the protein in GBM tissue samples, perhaps because it is a rather small and membrane bound protein (Fig. 7K).

### Circulating proteolytic product levels correlate with proteolytic activity in tumors

As described above for tissues, we also conducted a semi-specific analysis with the serum samples. We identified slightly more semi-specific than fully-tryptic peptides (4603 vs. 4501 - Fig. 8A). Interestingly, although we had observed a strong increase in inflammation markers, the fraction of semi-specific peptides remained unchanged following surgery for both primary and recurrent tumors (Fig. 8B). Hence, a detectable accumulation of proteolytic products might require sustained inflammation beyond the three days between surgery and postOP serum collection. When we compared preOP serum samples according to the cluster of their corresponding tissue, a significant increase of circulating semi-specific peptides paralleling the tissue’s proteolytic activity became apparent. This pattern was repeated in postOP samples, albeit without significance (Fig. 8C). Most detected cleavage products originated from canonical serum proteins, which might have come into contact with proteases from the inflamed tumor due to disruptions of the blood-brain barrier ^81^.

**Fig. 8.**
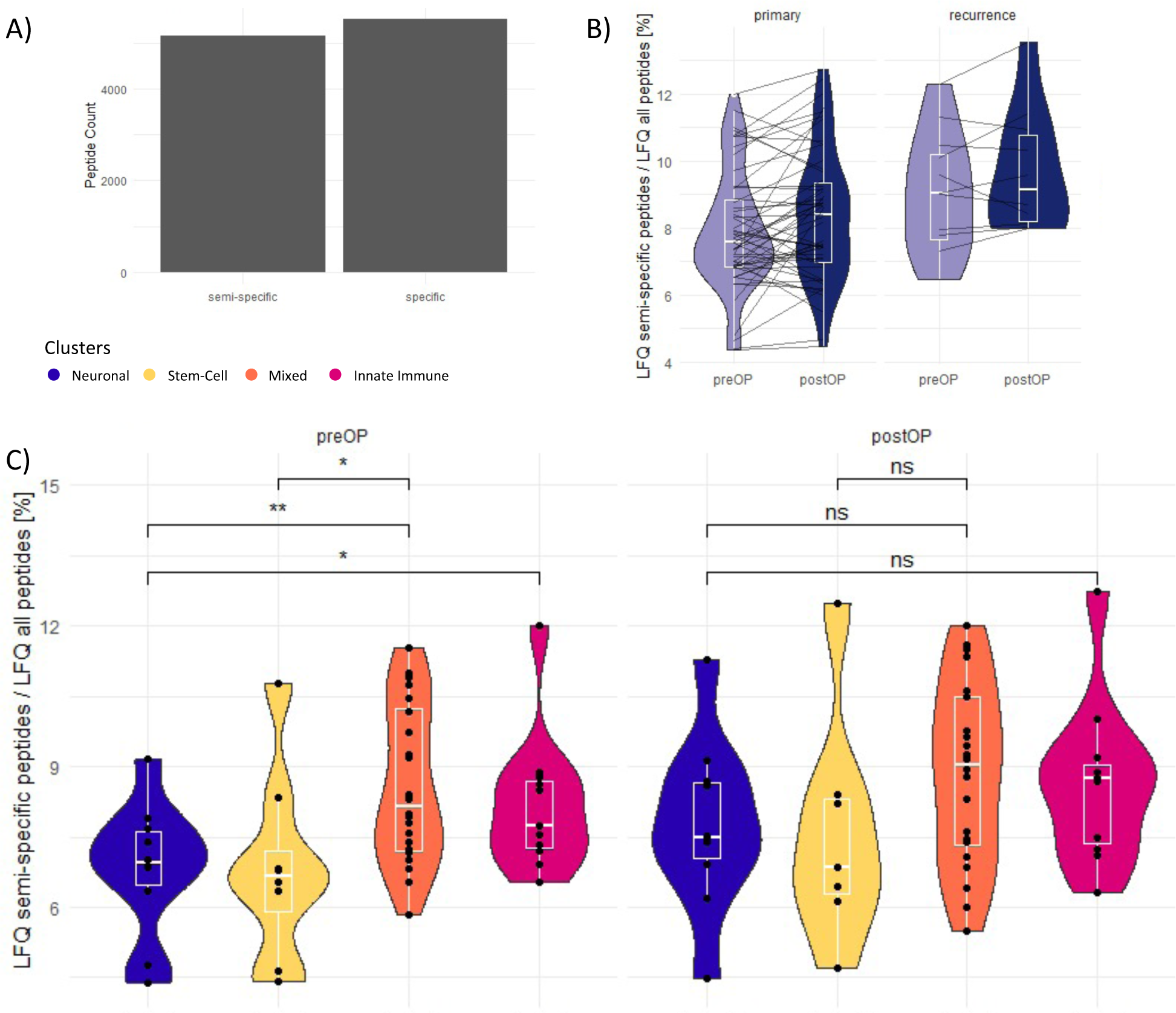
Semi-specific analysis of GBM serum. A) Identified semi-specific and fully tryptic peptides. B) Fraction of detected semi-specific peptides pre OP and post OP in primary and recurrent tumors. Matching samples are connected. C) Fraction of detected semi-specific peptides linked to the cluster of the corresponding primary tumor.

## Conclusion

In conclusion, we performed a comprehensive analysis of the proteomic landscape in matching tissue and serum samples from 55 GBM patients. We defined four distinct proteomic subtypes of primary GBM - neuronal, innate immunity, mixed, and stem-cell, which exhibited interconnected features. Our findings revealed the influence of immune infiltration and markers of neuronal development in shaping these proteomic subtypes, shedding light on the complex interplay between the tumor and its microenvironment. Further analyses, including missingness analysis and semi-specific proteolytic product identification, confirmed and expanded upon the biological motifs characterizing each cluster. Patients belonging to the neuronal cluster exhibited significantly longer survival than patients from the stem cell cluster. The innate immune cluster demonstrated high proteolytic activity which was aligned with its pronounced upregulation of immune-related proteins. In addition, our cohort contained five matching recurrences. Compared to their primary counterparts, these showed high proteomic plasticity with downregulation of immune and inflammatory proteins and upregulation of neuronal growth-related proteins. In the serum cohort, we identified potential circulating markers that could originate from the tumor tissue and serve as indicators of disease progression. PLA2G7 expression decreased significantly after primary and recurrence surgery, which was not the case in the MNG cohort. Additionally, a PAR2-peptide emerged as a candidate, exhibiting consistent changes after surgery in patients with inflamed tumors. Moreover, the abundance of circulating proteolytic products correlated to the proteolytic activity in the tumor and PAR2-peptide levels. Overall, our study underscores the intricate interplay between the GBM proteome, immune responses, and neurogenesis. The identified proteomic subtypes and potential circulating markers provide a foundation for further investigations aimed at understanding the underlying mechanisms of GBM progression and developing novel diagnostic and therapeutic strategies.

## Abbreviations

ABC: Ammonium Bicarbonate
AGC: Automatic Gain Control
ALB: Albumin
BCA: Bicinchonic Acid Assay
BLBP: Fatty acid-binding protein, brain
CAA: 2-Chloroacetamide
CD163: Scavenger receptor cysteine-rich type 1 protein M130
CPHM: Cox Proportional Hazards Model
CT: Computer Tomography
CTSG: Cathepsin G
DIA: Data Independent Acquisition
EDTA: Ethylenediaminetetraacetic acid
EGFR: Epidermal growth factor receptor
ENO2: Gamma-enolase
FCN1: Ficolin-1
FDR: False Discovery Rate
GBM: Glioblastoma Multiforme
GFAP: Glial fibrillary acidic protein
HCD: Higher-energy collisional dissociation
HEPES: 4-(2-hydroxyethyl)-1-piperazineethanesulfonic acid
HPLC: High Pressure Liquid Chromatography
HR: Hazard Ratio
HRAS: GTPase HRas
IDH: Isocitrate Dehydrogenase
IQR: Inter-Quartile Range
iRTs: Indexed Retention Time Standards
KRAS: GTPase KRas
L1CAM: Neural cell adhesion molecule L1
LC-MS/MS: Liquid-Chromatography Tandem Mass-Spectrometry
LFQ: Label-Free Quantification
LPA: Apolipoprotein (a)
LTQ: Linear Quadrupole Ion Trap Mass-Spectrometer
LUM: Lumican
LysC: Lysyl Endopeptidase C
LYZ: Lysozyme C
MAP2: Microtubule-associated protein 2
MARS: Multi Affinity Removal System
MNG: Meningioma
MRC1: Macrophage mannose receptor 1
MRI: Magnetic Resonance Imaging
mRNA: Messenger Ribonucleic Acid
mTOR: Mammalian Target of Rapamycine
NCAM1: Neural cell adhesion molecule 1
NES: Nestin
nLC: Nanoflow Liquid Chromatography
NOTCH1: Neurogenic locus notch homolog protein 1
NTRK2: BDNF/NT-3 growth factors receptor
NTRK3: NT-3 growth factor receptor
PAR2: Proteolysis Activated Receptor 2
PCA: Principal Component Analysis
PLS-DA: Partial-Least Squares Discriminant Analysis
PMSF: Phenylmethylsulfonyl Fluoride
postOP: Collection after Surgery
preOP: Collection before Surgery
PLA2G7: Platelet-activating factor acetylhydrolase
PTEN: Phosphatidylinositol 3,4,5-trisphosphate 3-phosphatase and dual-specificity protein phosphatase PTEN
RTN4: Reticulon-4
SDS: Sodium Dodecyl Sulfate
SOX10: Transcription factor SOX-10
SOX2: Transcription factor SOX-2
SP3: Single-Pot Solid-Phase-Enhanced Sample Preparation
SYP: Synaptophysin
SYT: Synaptotagmin-1
TCEP: Tris(2-carboxyethyl)phosphine hydrochloride
TFA: Trifluoroacetic acid
TMZ: Temozolomide
UMAP: Uniform Manifold Approximation and Projection for Dimension Reduction
VNN1: Pantetheinase
WHO: World Health Organization
ZEB1: Zinc finger E-box-binding homeobox 1

## Acknowledgements

This work has been supported by EPIC-XS, project number 823839, funded by the Horizon 2020 programme of the European Union.

The proteomics analyses were performed in the CRG/UPF Proteomics Unit which is part of the Spanish National Infrastructure for Omics Technologies (ICTS OmicsTech).

## Data Availability

Upon journal submission, the proteomic raw data will be accessible under restricted access for medical data protection reasons. For access before that, kindly contact the corresponding author at.

## Supplementary Data

- Table S1: Serum primary vs. recurrence
- Figure S1: GBM tissue cohort
- Figure S2: Clusters in primary tumor tissues
- Figure S3: Immune Markers
- Figure S4: UMAP and CoxBoost
- Figure S5: Primary vs. recurrent GBM tumor tissues
- Figure S6: Patient serum
- Figure S7: TCGA Expression Data

## Contributions

JWB and OS designed the project; AS, CN, AP, LC, AG, and JWB assembled the cohort and collected clinical data; TW, MH, and TF processed all samples; TW, GE, and ES performed the measurements; TW, MCC, and NP analyzed the data; TW, JWB, and OS wrote the manuscript.

