## Supplementary Tables and Figures for "Proteomic profiling of IDH-wildtype Glioblastoma Tissue and Serum uncovers prognostic Subtypes and Marker Candidates"

| | | full cohort (55 patients) | patient-matched prim-recurr samples (9 patients) | | | MNG control (26 patients) | |
|---|---|---|---|---|---|---|---|
| **Gene** | **UniProt ID** | **post1-pre1** | **post1-pre1** | **pre2-post1** | **post2-pre2** | **post-pre** | **Protein** |
| KIT | P10721 | -0,56 | -0,97 | 0,55 | -0,66 | (-0,33) | Mast/stem cell growth factor receptor Kit |
| C1QTNF3 | Q9BXJ4 | -0,55 | -0,83 | 0,72 | -1,04 | (-0,52) | Complement C1q tumor necrosis factor-related protein 3 |
| PLA2G7 | Q13093 | -0,61 | -0,81 | 0,80 | -0,69 | (-0,27) | Platelet-activating factor acetylhydrolase |
| CRTAC1 | Q9NQ79 | -0,62 | -0,79 | 0,71 | -0,49 | -0,55 | Cartilage acidic protein 1 |
| CNTN1 | Q12860 | -0,61 | -0,77 | 0,61 | -0,46 | -0,4 | Contactin-1 |
| CSF1R | P07333 | -0,52 | -0,77 | 0,68 | -0,52 | -0,46 | Macrophage colony-stimulating factor 1 receptor |
| AHSG | P02765 | -0,79 | -0,75 | 1,33 | -0,55 | -0,49 | Alpha-2-HS-glycoprotein |
| THBS4 | P35443 | -0,58 | -0,63 | 0,87 | -0,50 | -0,57 | Thrombospondin-4 |
| LUM | P51884 | -0,47 | -0,53 | 0,93 | -0,43 | (-0,23) | Lumican |
| CCDC66 | A2RUB6 | -0,55 | -0,53 | 0,46 | -0,22 | NA | Coiled-coil domain-containing protein 66 |
| ECM1 | Q16610 | -0,56 | -0,50 | 0,99 | -0,69 | -0,48 | Extracellular matrix protein 1 |
| PCOLCE | Q15113 | -0,55 | -0,48 | 0,89 | -0,97 | -0,46 | Procollagen C-endopeptidase enhancer 1 |
| BCHE | P06276 | -0,36 | -0,42 | 0,32 | -0,29 | -0,42 | Cholinesterase |
| COMP | P49747 | -0,72 | -0,42 | 0,84 | -0,72 | -0,58 | Cartilage oligomeric matrix protein |
| TPRA1 | Q86W33 | -0,28 | -0,39 | 0,48 | -0,23 | NA | Transmembrane protein adipocyte-associated 1 |
| ATRN | O75882 | -0,34 | -0,37 | 0,32 | -0,21 | -0,29 | Attractin |
| FCN3 | O75636 | -0,39 | -0,37 | 0,38 | -0,24 | (-0,31) | Ficolin-3 |
| CFB | P00751 | -0,31 | -0,34 | 0,40 | -0,25 | (-0,06) | Complement factor B |
| SERPINA7 | P05543 | -0,27 | -0,30 | 0,29 | -0,16 | (-0,22) | Thyroxine-binding globulin |
| AFM | P43652 | -0,3 | -0,19 | 0,37 | -0,26 | (-0,28) | Afamin |
| PROC | P04070 | 0,22 | 0,30 | -0,50 | 0,35 | (-0,14) | Vitamin K-dependent protein C |
| CWC22 | Q9HCG8 | 0,17 | 0,30 | -0,52 | 0,20 | NA | Pre-mRNA-splicing factor CWC22 homolog |
| TNC | P24821 | 0,55 | 0,32 | -1,21 | 0,84 | (-0,14) | Tenascin |
| AGT | P01019 | 0,36 | 0,33 | -1,21 | 0,71 | (0,23) | Angiotensinogen (Serpin A8) [Cleaved into: Angiotensin-1 |
| C5 | P01031 | 0,23 | 0,34 | -0,58 | 0,26 | (0,22) | Complement C5 |
| SERPING1 | P05155 | 0,27 | 0,37 | -0,56 | 0,28 | (0,23) | Plasma protease C1 inhibitor |
| GPX3 | P22352 | 0,43 | 0,38 | -1,12 | 0,66 | (0,25) | Glutathione peroxidase 3 |
| ARMCX5 | Q6P1M9 | 0,32 | 0,39 | -0,44 | 0,27 | NA | Armadillo repeat-containing X-linked protein 5 |
| MRC1 | P22897 | 0,54 | 0,42 | -0,73 | 0,46 | (0,30) | Macrophage mannose receptor 1 |
| PTPRF | P10586 | 0,45 | 0,45 | -0,83 | 0,28 | (0,21) | Receptor-type tyrosine-protein phosphatase F |
| HABP2 | Q14520 | 0,36 | 0,47 | -0,36 | 0,41 | (0,25) | Hyaluronan-binding protein 2 |
| VWF | P04275 | 0,59 | 0,66 | -1,33 | 0,77 | (0,36) | von Willebrand factor |
| COX8A | P10176 | 0,79 | 0,68 | -1,33 | 0,62 | NA | Cytochrome c oxidase subunit 8A, mitochondrial |
| GLE1 | Q53GS7 | 0,76 | 0,71 | -0,71 | 0,70 | NA | Nucleoporin GLE1 |
| SPANXN1 | Q5VSR9 | 0,82 | 0,74 | -1,35 | 0,78 | NA | Sperm protein associated with the nucleus on the X chromosome N1 |
| FSCB | Q5H9T9 | 0,85 | 0,77 | -1,39 | 0,87 | NA | Fibrous sheath CABYR-binding protein |
| ITGAL | P20701 | 0,86 | 0,85 | -1,05 | 0,63 | NA | Integrin alpha-L |
| SERPINA3 | P01011 | 0,84 | 0,86 | -1,27 | 0,72 | 0,61 | Alpha-1-antichymotrypsin |
| ALB | P02768 | 0,96 | 0,90 | -2,26 | 2,93 | (0,14) | Albumin |
| HP | P00738 | 1,48 | 1,52 | -2,42 | 1,76 | 1,37 | Haptoglobin |
| CLLU1-AS1 | Q5K130 | 1,43 | 2,46 | -2,19 | 1,49 | NA | Putative uncharacterized protein CLLU1-AS1 |

**Table S1 Serum primary vs. recurrence.** Result of serum differential expression analysis. All log2-fold changes given without parentheses were significant p≤0.05. Coloring indicates how the expression change relates to preOP-postOP expression changes in MNG cohort. Blue: similar change in MNG cohort. White: not detected in MNG cohort. Yellow: Unchanged in MNG cohort.

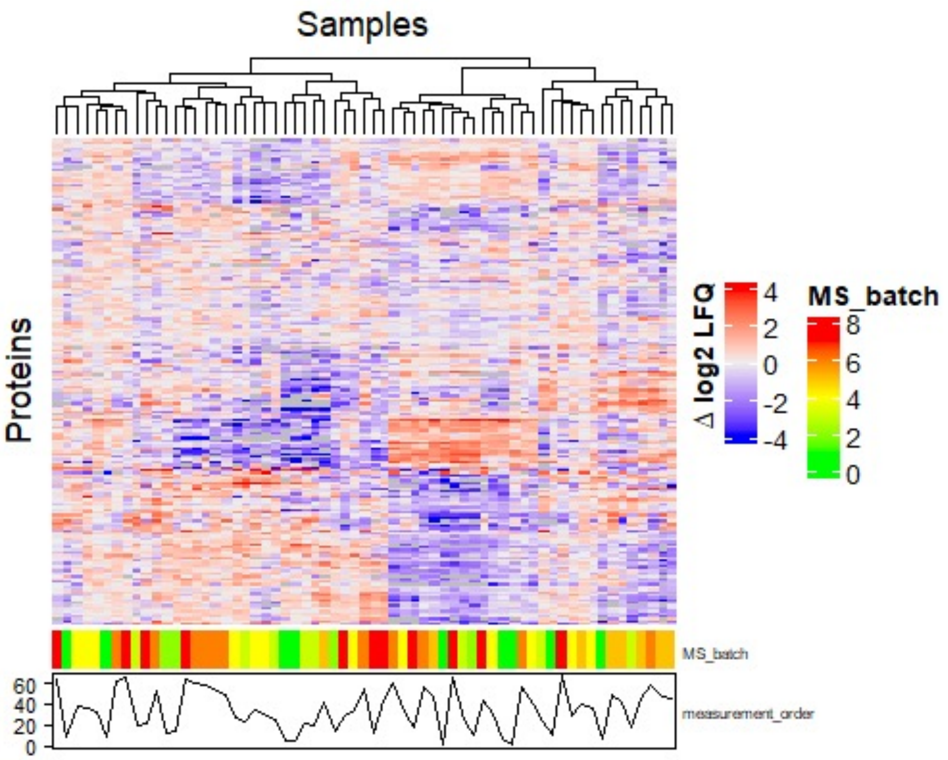


**Fig. S1 GBM tissue cohort.** Heatmap of the GBM tissue cohort. Measurement batches and order are given below.

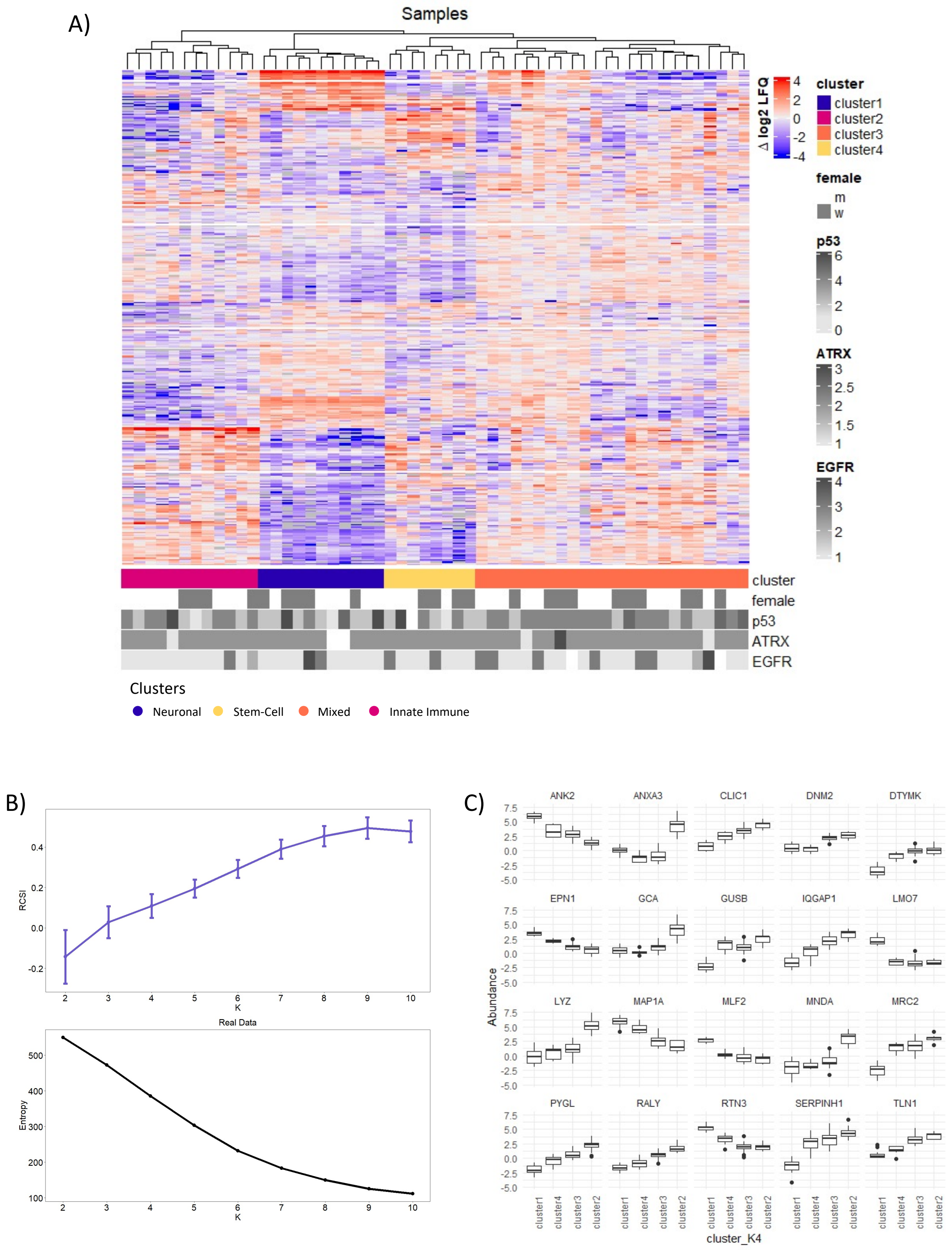


**Fig. S2 Clusters in primary tumor tissues.** A) Heatmap of the GBM tissue cohort. Hierarchical clusters including clinical variables are shown below. B) Monte-Carlo simulation results for optimal number of hierarchical clusters. C) Boxplots of differential expression analysis between clusters showing the top 20 most differentially abundant proteins.

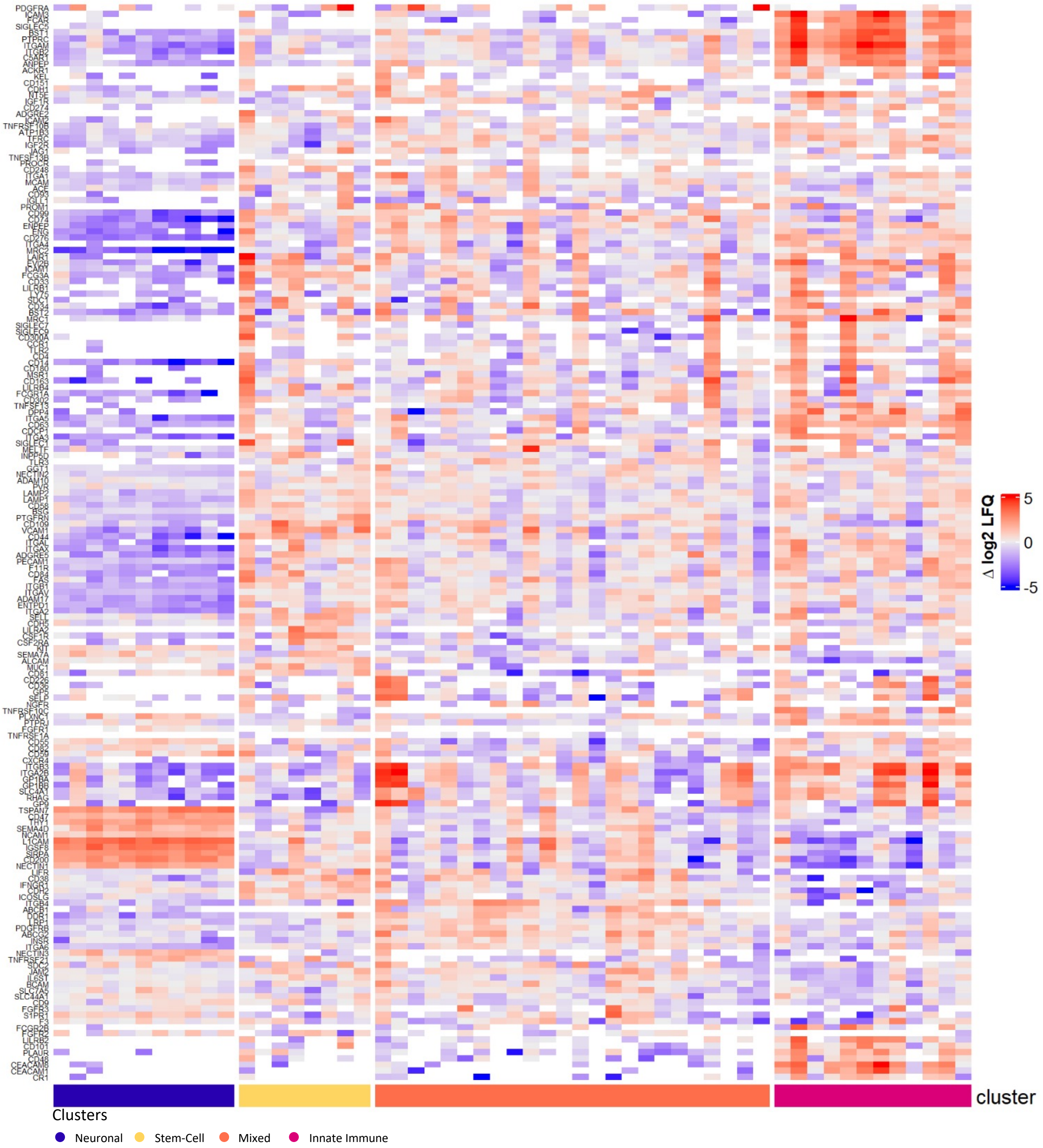


**Fig. S3 Immune Markers.** Heatmap of all identified immune marker proteins.

A)

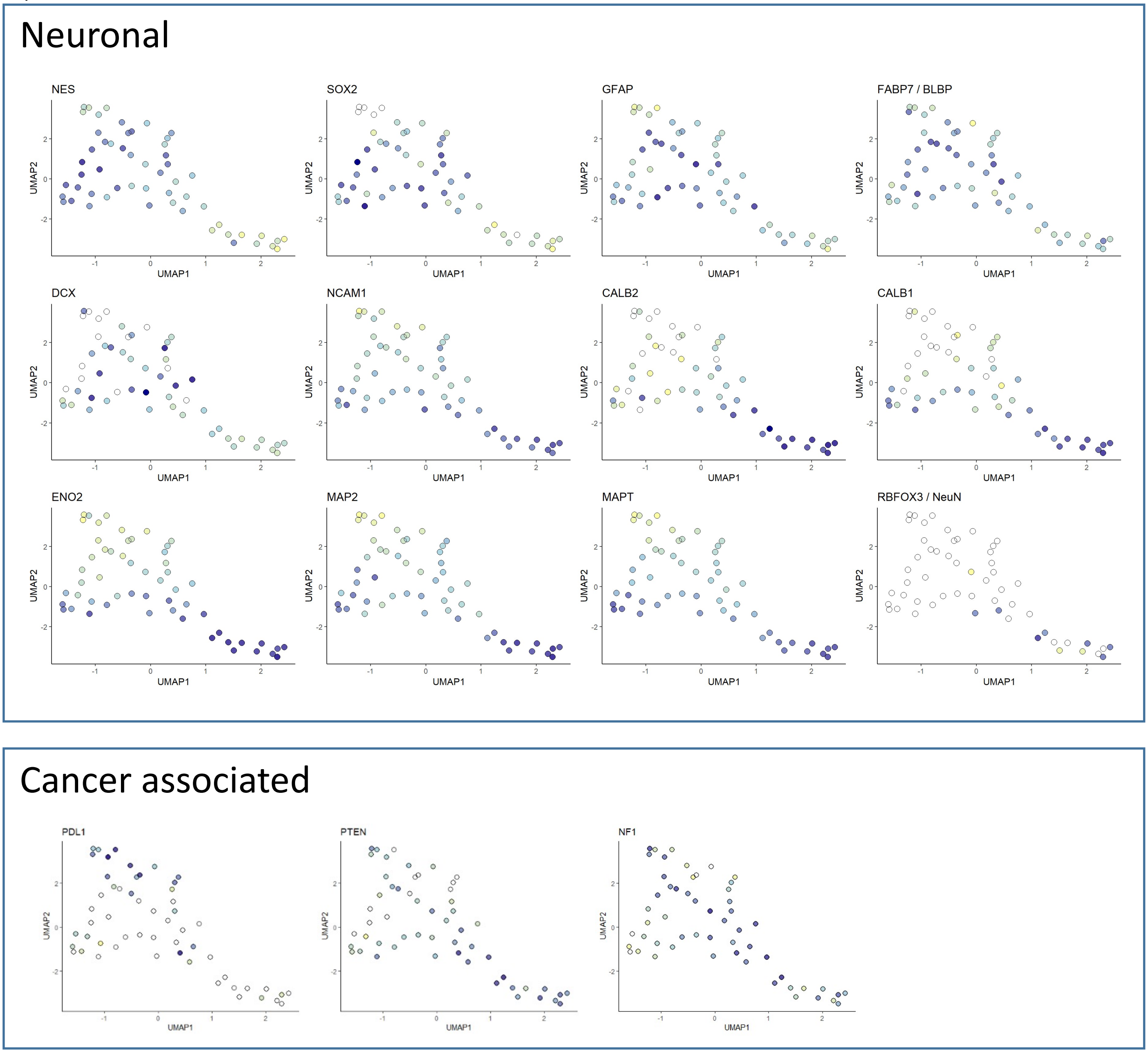


B)

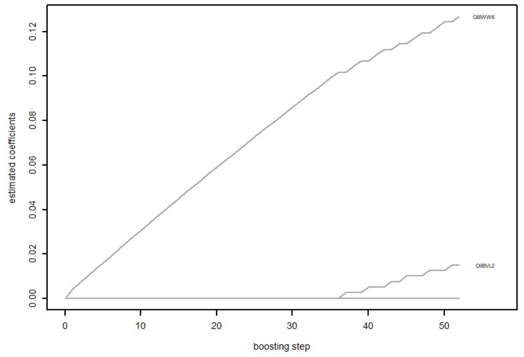


**Fig. S4 UMAP and CoxBoost.** A) UMAPs of different proteins of interest including immune cell markers, neurogenesis and differentiation markers, and proteins associated with cancer. B) CoxBoost result.

A)

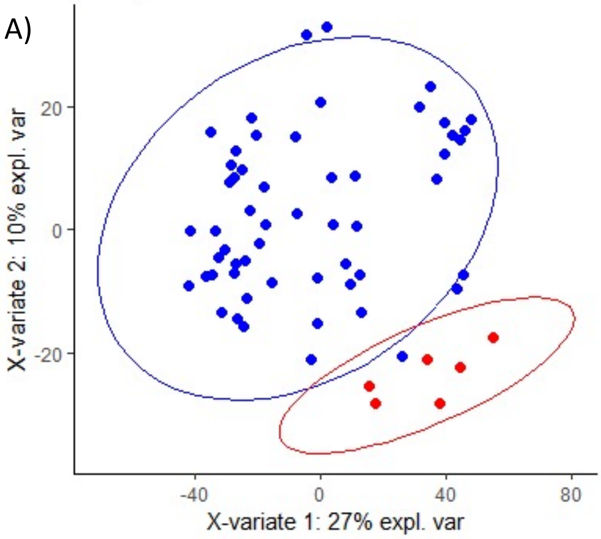


B)

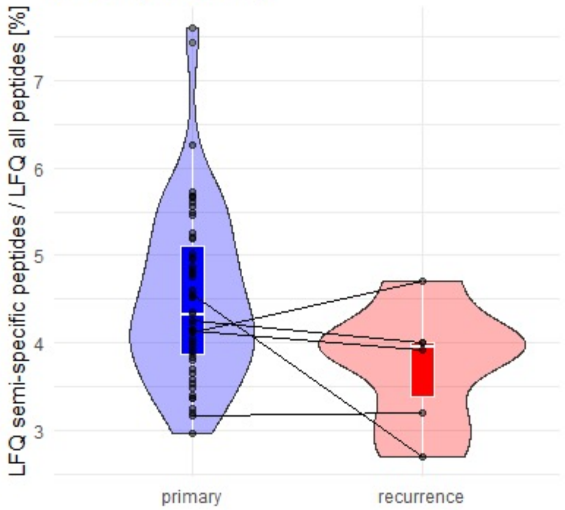


**Fig. S5 Primary vs. recurrent GBM tumor tissues.** Primary tumors are shown in blue, recurrent tumors in red. A) PLS-DA of all GBM tissues. B) Fraction of detected semi-specific peptides between primary and recurrent tumors. Matching sample pairs are connected.

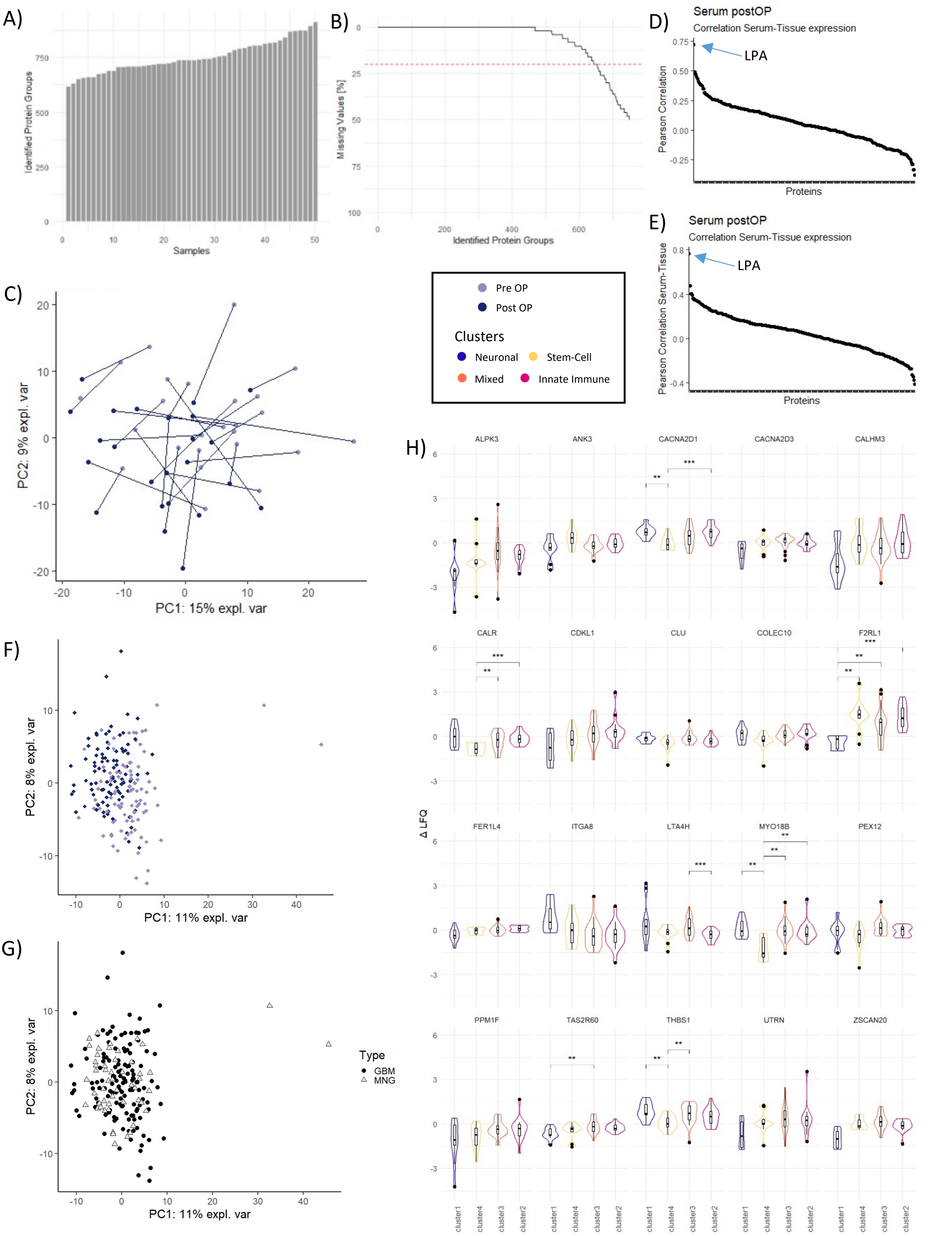


**Fig. S6 Patient serum.** A) Identified Proteins in MNG cohort. B) Missingness in MNG cohort. C) PCA of MNG cohort. Matching samples are connected. D & E) Pearson correlation coefficients between proteins identified in GBM serum and tissue for pre OP (D) and post OP (E) samples. F & G) PCAs of GBM and MNG serum cohorts merged via ComBAT. H) Violin plots of differential expression analysis between clusters showing the top 20 most differential protein abundance changes between pre OP and post OP serum

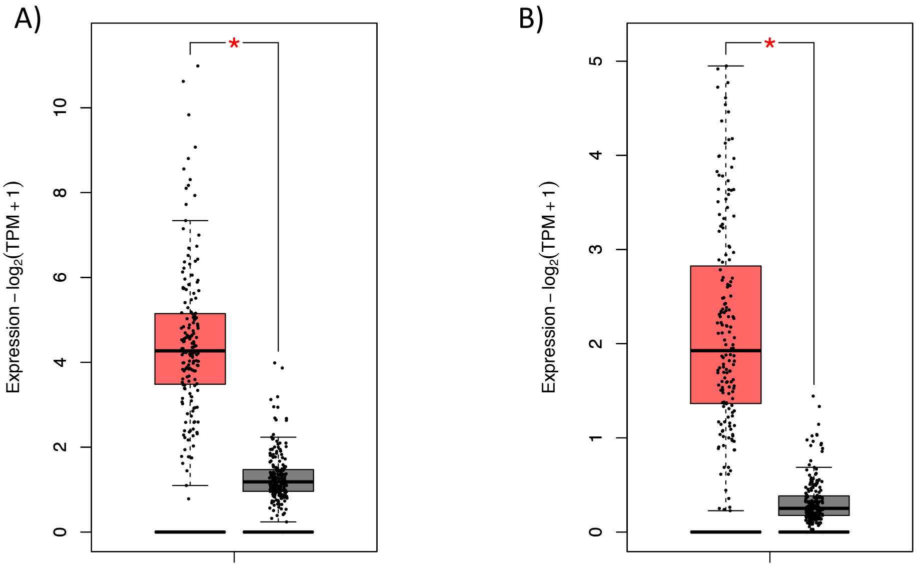


**Fig. S7 TCGA Expression Data.** Gene expression in normal brain (NB) tissue and GBM patient samples given in Transcripts per Million (TPM). Data were obtained from TCGA and GTEX data (www. gepia2.cancer-pku.cn) on n = 207 (NB, grey) and n = 163 (GBM, red) individuals. Expression difference is significant with $p < 0.01$. A) *LUM*/LUM. B) *F2RL1*/PAR2.
